# The *SLC1A1*/EAAT3 Dicarboxylic Amino Acid Transporter is an Epigenetically Dysregulated Nutrient Carrier that Sustains Oncogenic Metabolic Programs

**DOI:** 10.1101/2023.09.04.556240

**Authors:** Treg Grubb, Jesminara Khatun, Sayed Matar, Fatme Ghandour, Noah Dubasik, Carleigh Salem, David A. Orlando, Matthew G. Guenther, Steven R. Martinez, Pooneh Koochaki, Jesse A. Coker, Cerise Tang, Eduard Reznik, Ritesh R. Kotecha, A. Ari Hakimi, Nour Abdallah, Christopher J. Weight, Toni K. Choueiri, John M. Asara, Shaun R. Stauffer, Sabina Signoretti, William G. Kaelin, Abhishek A. Chakraborty

**Affiliations:** Genitourinary Malignancies Research Center, Department of Cancer Biology, Lerner Research Institute, Cleveland Clinic, Cleveland, OH, USA; Department of Pathology, Brigham and Women’s Hospital, Boston, MA, USA; Syros Pharmaceuticals, Inc, Cambridge, MA, USA; Center for Therapeutics Discovery, Cleveland Clinic, Cleveland, OH, USA; Department of Surgery, Memorial Sloan Kettering Cancer Center, New York, NY, USA; Department of Urology, Glickman Urological and Kidney Institute, Cleveland, OH; Department of Medical Oncology, Lank Center for Genitourinary Oncology, Dana-Farber Cancer Institute, Brigham and Women’s Hospital, and Harvard Medical School, Boston, MA, USA; Mass Spectrometry Core, Beth Israel Deaconess Medical Center, Department of Medicine, Harvard Medical School, Boston, MA, USA; Department of Medical Oncology, Dana-Farber Cancer Institute and Brigham and Women’s Hospital, Harvard Medical School, Boston, MA 02215, USA; Howard Hughes Medical Institute, Chevy Chase, MD 20815, USA; Case Comprehensive Cancer Center, Case Western Reserve University, Cleveland, OH 44106, USA

## Abstract

Inactivation of pVHL tumor suppressor in clear cell Renal Cell Carcinoma (ccRCC) increases the abundance of Histone H3 lysine 27 acetylation (H3K27ac). We hypothesized that H3K27ac, a marker of transcriptional activation, drives the expression of critical oncogenes in ccRCC. Using H3K27ac ChIP-Seq; RNA-Seq; an *in vivo* positive selection screen; cell-based functional studies; and clinical validations; here, we report the identification of the SLC1A1/EAAT3 aspartate (Asp) and glutamate (Glu) transporter as a ccRCC oncogene. pVHL loss promotes SLC1A1 expression in a HIF-independent manner. Importantly, SLC1A1 inactivation depletes Asp/Glu-derived metabolites, impedes ccRCC growth both *in vitro* and *in vivo*, and sensitizes ccRCCs to metabolic therapeutics (e.g., glutaminase blockers). Finally, in human ccRCC biospecimens, higher SLC1A1 expression is associated with metastatic disease and clusters with elevated expression of other solute carriers, but not HIF/Hypoxia pathways. Altogether, our studies identify a HIF-independent metabolic hub in ccRCC and credential SLC1A1 as an actionable ccRCC oncogene.

**STATEMENT OF SIGNIFICANCE:** Targeting chronic HIF activation underlies many therapeutic strategies in ccRCC; but, unfortunately, is not curative. SLC1A1, instead, represents a HIF-independent ccRCC dependency, which is targetable alone and together with other antimetabolites, such as glutaminase inhibitors. These observations identify an actionable metabolic program that functions independent of HIF in ccRCC.

## INTRODUCTION

Kidney cancer, or Renal Cell Carcinoma (RCC), is among the ten most common forms of cancer in both men and women (1). The three major sub-types of RCC include clear cell RCCs (ccRCCs), which typically present with loss of the von Hippel-Lindau tumor suppressor protein (pVHL); and the papillary (pRCC) and chromophobe (chRCC) cancers, which are typically pVHL-proficient (2).

pVHL functions as the substrate recognition subunit in a Cullin2-dependent ubiquitin ligase (E3) complex (3). Perhaps, the best-known cellular function of the pVHL E3 ligase is the oxygen-dependent proteolysis of the α-subunit of the Hypoxia Inducible Factor (HIFα) transcription factors (4, 5). Therefore, pVHL-deficient ccRCCs exhibit chronic activation of HIFα and hyperactivated HIF transcription programs, particularly those controlled by HIF2α, drive ccRCC oncogenesis (6, 7). These observations have justified the development of pharmacological HIF2α inhibitors (8, 9), one of which is now FDA approved for the treatment of ccRCC in VHL syndrome patients who harbor germline mutations in pVHL (10).

Importantly, patients with sporadic ccRCC also routinely harbor truncal loss-of-function mutations in pVHL (11, 12); however, these patients have shown lower responses to HIF2α blockade. This suggests that either greater genetic heterogeneity and/or extensive prior therapy in the clinical trial cohorts diminishes HIF2α dependence in sporadic tumors. Identifying pVHL-dependent – but HIF-independent – oncogenic programs are thus of immense therapeutic value.

Chromatin dysregulation is a hallmark feature of ccRCC and occurs via both HIF-dependent and independent mechanisms (13). HIF-dependent mechanisms include the transcriptional activation of several JumonjiC-family histone demethylases (KDMs) (14), whereas HIF-independent mechanisms include the genetic inactivation of SWI/SNF-family chromatin modifiers, such as *PBRM1* (15); or inactivation of histone modifiers, such as *SETD2*, *MLL3*, *KDM5C*, and *KDM6A* (16). These changes likely select for oncogenic epigenetic programs in ccRCC, but also render these tumors dependent on the compensatory function of other chromatin/histone modifiers (17). Epigenetic dysregulation, therefore, represents an exploitable vulnerability in kidney cancer.

To begin exploring the oncogenic importance of epigenetic dysfunction, we previously profiled changes in histone modification patterns in pVHL-deficient ccRCCs (versus the pVHL-proficient pRCCs and chRCCs) and observed that ccRCCs have markedly elevated levels of acetylated lysine 27 on Histone H3 (H3K27ac) (17). H3K27ac is associated with transcriptional activation and its accumulation at cis-regulatory elements (e.g., promoters and enhancers/super-enhancers) marks key oncogenes and regulators of cellular identity in many cancers (18, 19). We hypothesized that elevated H3K27ac in pVHL-deficient ccRCCs, likewise, marks critical oncogenes that sustain tumorigenic programs in kidney cancer. Identifying these epigenetic targets could yield unexplored strategies to control the spread of kidney cancer.

## RESULTS

### Dysregulated H3K27ac Patterns Identify Candidate ccRCC Oncogenes

Cell lines established from human patient specimens have facilitated the pre-clinical discovery of actionable targets in ccRCC for many decades. Reintroduction of pVHL does not prominently impact the *in vitro* growth of pVHL-deficient ccRCC cell lines under standard culture conditions. Therefore, lentivirally transducing ccRCC cells to restore expression of pVHL (or empty vector) has allowed us to previously generate isogenic pairs of a number of ccRCC cell lines. Using one representative isogenic pair (UMRC-2 cells, either with or without pVHL expression), we profiled for pVHL-dependent changes in H3K27ac deposition. Using ChIP-Seq analysis, we annotated ∼8,400 peaks that were differentially marked by H3K27ac between pVHL-deficient (VEC) versus pVHL-proficient cells (Fig. 1A).

**Figure 1.**
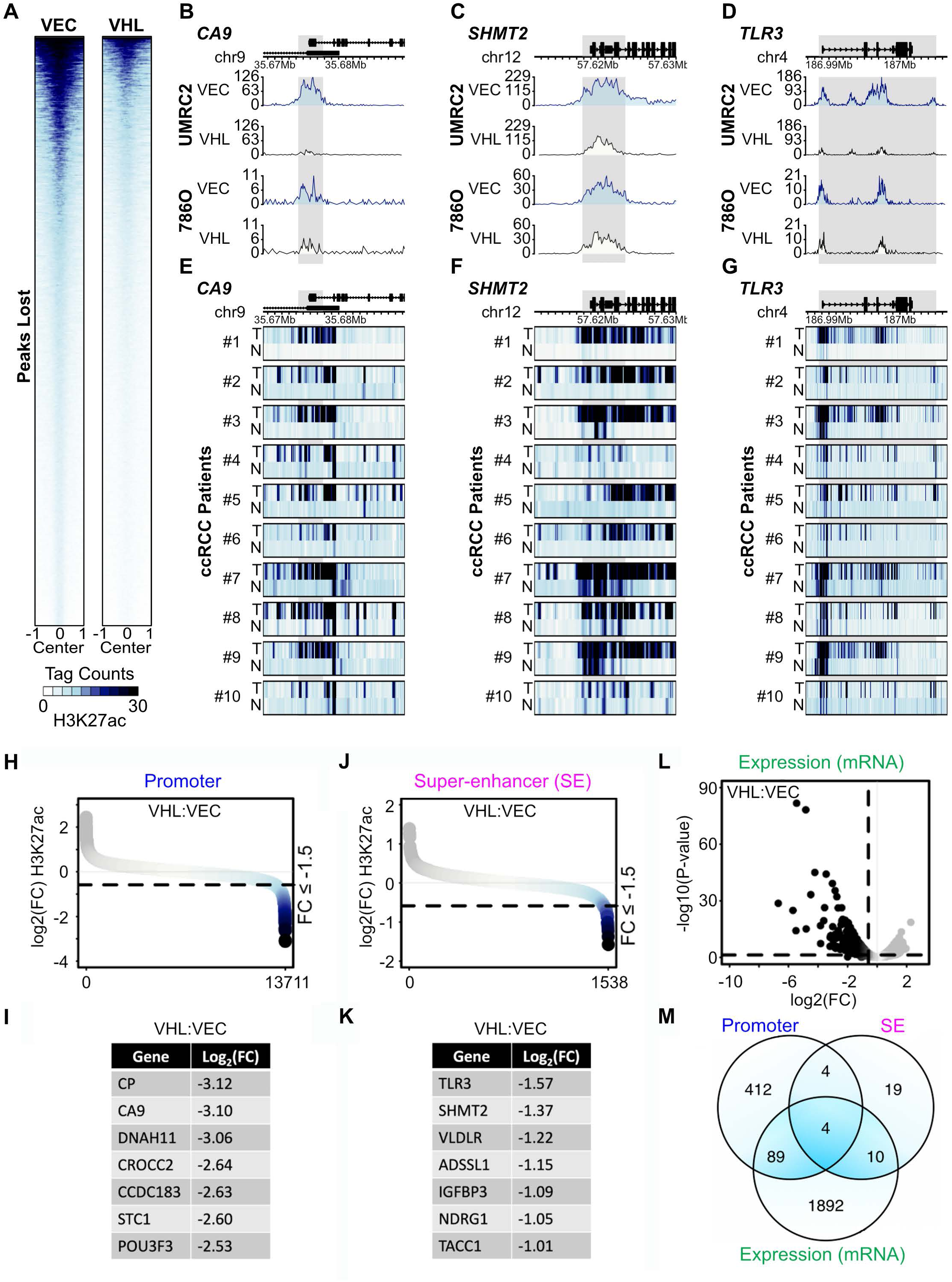
pVHL-dependent Epigenetic Dysregulation Identifies Candidate Oncogenes. (**A**) Heatmap representing the abundance of H3K27ac peaks scored as decreased (≥1.5 fold) in UMRC-2 cells that were lentivirally transduced to express pVHL (VHL) or empty vector (VEC). (**B** to **D**) H3K27ac signal at thwe *CA9* (**B**), *SHMT2* (**C**), or *TLR3* (**D**) proximal regions in pVHL-proficient (VHL) and pVHL-deficient (VEC) versions of two ccRCC cell lines (UMRC-2 and 786-O). (**E** to **G**) H3K27ac signal at the *CA9* (**E**), *SHMT2* (**F**), or *TLR3* (**G**) proximal regions, as described in (B) to (D), in human ccRCC tumors versus their paired adjacent normal tissue. (**H** to **K**) H3K27ac signal plotted as a log_2_[fold-change] ratio between pVHL-proficient versus pVHL-deficient UMRC-2 cells at promoter regions (**H** and **I**) or enhancers/super-enhancers (**J** and **K**). (**L**) Volcano plot of RNA-Seq data showing differentially expressed genes in cells described in (H to K). (**M**) Venn diagram indicating the overlap between pVHL-dependent changes in H3K27ac in the promoter and enhancer/super-enhancers (SE) and changes in mRNA abundance to generate a candidate gene list. For Promoter and SE datasets a FC of ≥1.5 was used as a cut-off, whereas for the mRNA dataset a FC of ≥2.0 was used as a cut-off.

We found that the H3K27ac peaks in pVHL-deficient UMRC-2 cells overlapped closely with a second representative ccRCC cell line (786-O). Importantly, as expected, the genes harboring elevated H3K27ac in the pVHL-deficient cells included many notable HIF target genes (e.g., *CA9* and *EGLN3*) and previously characterized ccRCC oncogenes [e.g., *SHMT2* (20, 21) and *ZHX2* (22)], but also included genes that are *not* presently associated with ccRCC (e.g., *TLR3* and *SLC28A1*) (Fig. 1B to D; Supplementary Table S1). Using a previously published dataset (23), we next compared H3K27ac deposition in human renal tumors (versus adjacent normal tissue). This comparison showed that many of the genomic regions marked by elevated H3K27ac in pVHL-deficient cells were, likewise, decorated with higher H3K27ac in the ccRCC tumors compared to their adjacent normal tissue (Fig. 1E to G; Supplementary Table S1), thus validating the physiological relevance of our cell line generated dataset. Together, these observations justified further studies on the functional importance of the relatively understudied H3K27ac marked loci.

To address functional relevance, we characterized the transcriptional output of the H3K27ac-marked genomic loci. By comparing H3K27ac at annotated cis-regulatory elements, including promoters (Fig. 1H and I) and enhancers/super-enhancers (Fig. 1J and K) with pVHL-dependent transcriptional changes determined by RNA-Seq (Fig. 1L), we established a candidate list of ∼100 genes that: (**1**) had higher H3K27ac in pVHL-deficient cells; (**2**) exhibited higher mRNA levels upon pVHL inactivation; and (**3**) encoded druggable gene products, such as cell-surface proteins, enzymes, etc. (Fig. 1M; Supplementary Table S2).

### Functional In Vivo Screens Score SLC1A1 as an Oncogenic Driver in ccRCC

pVHL-proficient cells, unlike their pVHL-deficient counterparts, do not efficiently form tumors *in vivo*. To functionally interrogate the importance of the epigenetically dysregulated candidate genes, we performed an *in vivo* positive selection (up) screen and sought to identify genes whose expression was sufficient to confer tumorigenic potential onto otherwise poorly tumorigenic pVHL-proficient cells. We manually curated an ORF library corresponding to our (∼100) candidate genes and included additional miscellaneous controls, such as BFP, GFP, and HcRed (Supplementary Table S2). The individual clones were procured from the Broad Institute in the pLX317-backbone, which has a unique 20 nucleotide barcode associated with each gene. We generated an equimolar pool of the individual clones, packaged this library into lentiviral particles, and transduced pVHL-proficient versions of UMRC-2 cells at low MOI. Transduced cells were expanded, a small aliquot (∼10%) was harvested and set aside to measure representation at the assay start (T0), and 5e6 cells of the remaining cells inoculated subcutaneously (flank xenografts) into immunodeficient NCR^nu/nu^ mice (Fig. 2A).

**Figure 2.**
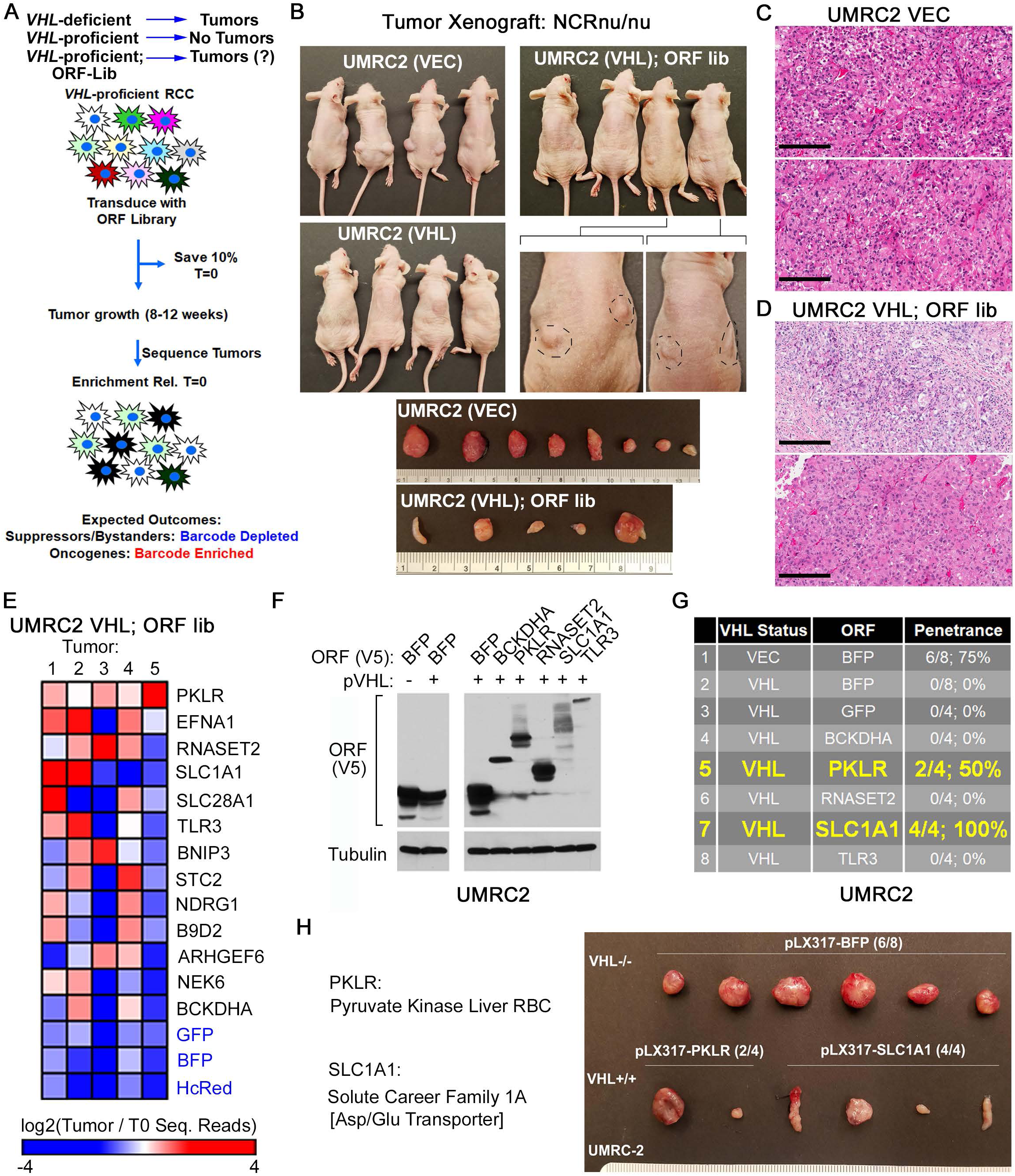
An In Vivo Positive Selection Screen Identifies SLC1A1 as a ccRCC Oncogene. (**A**) Schema of the *in vivo* positive selection screen to identify genes whose expression is sufficient to confer tumorigenic potential onto pVHL-proficient ccRCCs. (**B** to **D**) Photographs showing tumor growth (**B**) and H&E stained sections (**C** and **D**) of the indicated versions of the UMRC-2 cells that were inoculated sub-cutaneously to form flank xenografts in NCR^nu/nu^ mice. Primary endpoint for pVHL-deficient (VEC) UMRC-2 cells was reached at 8 weeks post injection and for the pVHL-proficient cells expressing the ORF library (VHL; ORF lib) at 12 weeks post injection. (**E**) Heatmap depicting the changes in relative abundance of barcodes representing the indicated genes, as measured by next-gen sequencing from genomic DNA harvested from the pVHL-proficient versions of the UMRC-2 cells that were lentivirally transduced with the ORF library (VHL; ORF lib). (**F** to **H**) Immunoblot analysis (**F**) and sub-cutaneous tumor growth (**G** and **H**) measured in pVHL-proficient versions of UMRC-2 cells lentivirally transduced to express the indicated genes and then inoculated sub-cutaneously into flanks of NCR^nu/nu^ mice.

As expected, our positive control pVHL-deficient UMRC-2 cells (VEC) formed tumors at 100% penetrance (8/8 injections), whereas pVHL-proficient UMRC-2 cells (VHL) failed to form tumors (0/8 injections) (Fig. 2B). Interestingly, introduction of the ORF-library into pVHL-proficient cells led to tumor growth in 5/8 injections (∼60% penetrance) (Fig. 2B to D). Tumors from the ORF-library transduced cells were harvested to measure ORF representation at the assay endpoint (Tend). Genomic DNA from the T0 and Tend samples was used to PCR amplify the pLX317 barcode region and compared using next-gen sequencing.

We anticipated that barcodes representing oncogenes would be enriched in tumors, whereas barcodes representing either neutral bystanders or tumor suppressors would be depleted. To account for biological differences, we chose to pursue candidates that scored as enriched in at least two of the five tumors. While our negative controls (e.g., BFP, GFP, and HcRed) were depleted in all five tumors, we identified eleven enriched candidate genes, many of which were known targets of HIF, such as *BNIP3* and *NDRG1* (Fig. 2E). We prioritized genes that were not canonical HIF-targets and in secondary validation experiments generated pVHL-proficient UMRC-2 and 786-O cells individually expressing these candidate genes (Fig. 2F; Supplementary Fig. S1A). *PKLR* expression promoted tumor growth at 50% penetrance (2/4 injections) in pVHL-proficient UMRC-2 cells, but not in 786-O cells. However, in both 786-O and UMRC-2 cells, reintroduction of *SLC1A1* conferred tumorigenic potential, albeit at 50% penetrance in 786-O cells (2/4 injections) and 100% penetrance in UMRC-2 cells (4/4 injections) (Fig. 2G and H; Supplementary Fig. S1B and S1C).

### SLC1A1 Function is Necessary for the Survival of pVHL-deficient ccRCCs

*SLC1A1* encodes a dicarboxylic amino acid transporter, also called Excitatory Amino Acid Transporter 3 (EAAT3), which drives the cellular uptake of Asp and Glu (and to a lesser extent Cys) (24, 25). Despite these amino acids being ‘non-essential’ under normal physiological conditions, we reasoned that they likely provide essential carbon skeletons to support the dysregulated biosynthetic needs of transformed pVHL-deficient ccRCCs. Consistent with this idea, CRISPR/Cas9-mediated inactivation of *SLC1A1* using three different sgRNAs, but not non-targeting controls, showed that SLC1A1 was necessary for survival in three human pVHL-deficient ccRCC lines (Fig. 3A and B).

**Figure 3.**
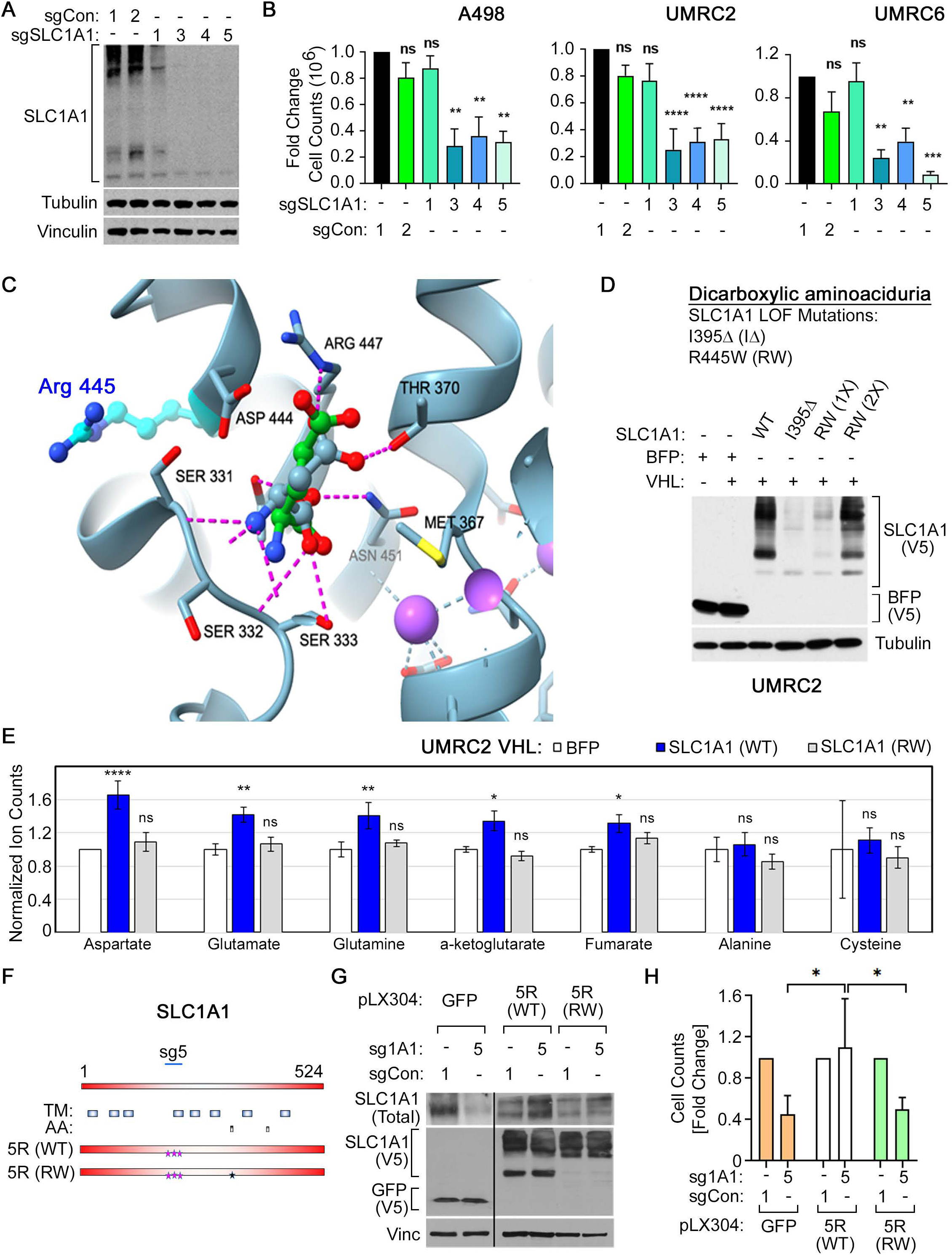
SLC1A1’s Metabolic Function is Necessary to Support ccRCC Growth. (**A** and **B**) Immunoblot analysis in UMRC-2 cells (**A**) and normalized cell numbers, relative to control sgRNA #1 (sgCon 1), as measured using the ViCell, in the indicated cell lines that were lentivirally transduced to express sgRNAs targeting SLC1A1 or non-targeting controls (sgCon), as indicated. Counts were compared using one-way ANOVA with Dunnett’s multiple comparison test, n=3, **p<0.01; ***p<0.001; ****p<0.0001; ns = non-significant. Cell counts in (**B**) for A-498 and UMRC-6 cells were done 7 days post selection and for UMRC-2 cells at 14 days post selection. (**C**) Structural modelling of Asp and Glu into the SLC1A1 ligand-binding pocket. (**D**) Immunoblot analysis measuring expression of two loss-of-function SLC1A1 mutants, deletion of Ile395 (I395Δ) and Arg445 ->Trp [R445W (RW)] or BFP as a miscellaneous control, expressed in pVHL-deficient or proficient versions of UMRC-2 cells, as indicated. (**E**) Normalized abundance, relative to total ion count, of the indicated metabolites, measured using LC-MS/MS from pVHL-proficient versions of UMRC-2 cells that were transduced with the indicated forms of SLC1A1 or a miscellaneous BFP control, as indicated. Ion counts were compared using two-way ANOVA (BFP vs WT and BFP vs RW) with Dunnett’s multiple comparison test, n=3, *p<0.05; **p<0.01; ****p<0.0001; ns = non-significant. (**F**) Schema showing SLC1A1 sg5 (sg1A1#5)-resistant versions of wild-type [5R (WT)] and RW mutant [5R (RW)] SLC1A1. Pink stars mark silent-mutations in the sgRNA-recognition sequence and the blue star marks the R445W mutation. (**G** and **H**) Immunoblot analysis (**G**) and cell counts measured using the ViCell (**H**) of UMRC-2 cells that were first lentivirally transduced to express the indicated forms of SLC1A1 (or GFP control) in the pLX304 backbone and then transduced to express either the sgRNA targeting SLC1A1 or a non-targeting control (sgCon). Immunoblot analysis (**G**) was done three days post-selection in Puromycin (2 µg/mL) and cell numbers were measured fourteen days post selection. Counts in the sg5-expressing arms in (H) were compared using Student’s t-test with Holm-Sidak correction for multiple testing, n=2, *p<0.05. Immunoblots in (**G**) were run on the same gel; however, because of the significantly greater expression of GFP, lower exposures of the first two lanes are presented alongside exposures with similar intensities for WT and RW SLC1A1.

We then modeled the ligand-binding pocket of SLC1A1 with its canonical ligands, Asp and Glu, and found key residues in SLC1A1 that directly interacted with these amino acids (e.g., Ser331-333, Asp444, and Arg447) (Fig. 3C). Our model closely resembles the recently published Cryo-EM structure of SLC1A1 (26). We compared our model with SLC1A1 loss-of-function mutations, which have been previously described in a human pathology called Dicarboxylic Aminoaciduria, such as Ile395Δ (I395Δ) and Arg445->Trp [R445W (RW)] (27). Interestingly, although not directly involved in ligand binding, we identified Arg445 as a residue that likely stabilizes the electrostatic interactions in the ligand-binding pocket (Fig. 3C).

Reintroduction of wild-type SLC1A1 into pVHL-proficient cells increased intracellular levels of Asp, Glu, and their downstream TCA derivatives, but did not impact pools of unrelated amino acids, such as Ala or Cys (Fig. 3D and E). These metabolic changes were specific and did not occur upon expression of BFP (control) or the functionally-dead RW variant. Finally, expressing a sgRNA-resistant version of wild-type SLC1A1 [5R (WT)], but not the RW mutant [5R (RW)], rescued the fitness defects associated with *SLC1A1* inactivation in pVHL-deficient cells (Fig. 3F to H). Altogether, these studies showed that the effects of the SLC1A1 sgRNAs were ‘on-target’ and suggested that the canonical function of SLC1A1 as an Asp/Glu importer was necessary for its oncogenic function in ccRCC.

### pVHL Inactivation Induces SLC1A1 Expression in a HIF-independent Manner

The EglN prolyl hydroxylases catalyze oxygen-dependent prolyl hydroxylation at two key proline residues on HIFα. Hydroxylated HIFα is then recognized by the pVHL ubiquitin ligase and targeted for ubiquitin-dependent proteolysis (28). Inactivation of the EglNs or of pVHL thus leads to stabilization of HIFα. Because chronic HIF activation is a prominent regulator of the transcriptional changes driven by pVHL loss, we addressed if the transcriptional induction of *SLC1A1* in pVHL-deficient cells was HIF-dependent.

We first confirmed that H3K27ac marking at the *SLC1A1* locus was diminished upon pVHL expression in the isogenic ccRCC cell lines and resembled the effects of pVHL introduction on H3K27ac at canonical HIF target genes, such as *EGLN3* and *NDRG1* (Fig. 4A). Interestingly, analysis of the human tumor dataset (23) confirmed increased H3K27ac marking at the *SLC1A1* locus in a subset of tumors versus the normal adjacent tissue (e.g., #1, #2, #3, #5, #6, and #8). Surprisingly, we noted stronger H3K27ac marking in some normal tissues (e.g., #7, #9, and #10) (Fig. 4B). We reasoned this H3K27ac marking in normal tissue represents the basal SLC1A1 expression to perform its canonical role in dicarboxyclic amino acid uptake. Lastly, using both western blotting and realtime-qPCR analysis, we validated our primary observations in three human ccRCC cell lines (UMRC-2, UMRC-6, and 786-O), and found that reintroduction of pVHL decreased both mRNA and protein level expression of SLC1A1 (Fig. 4C and D; Supplementary Fig. S2A to S2D).

**Figure 4.**
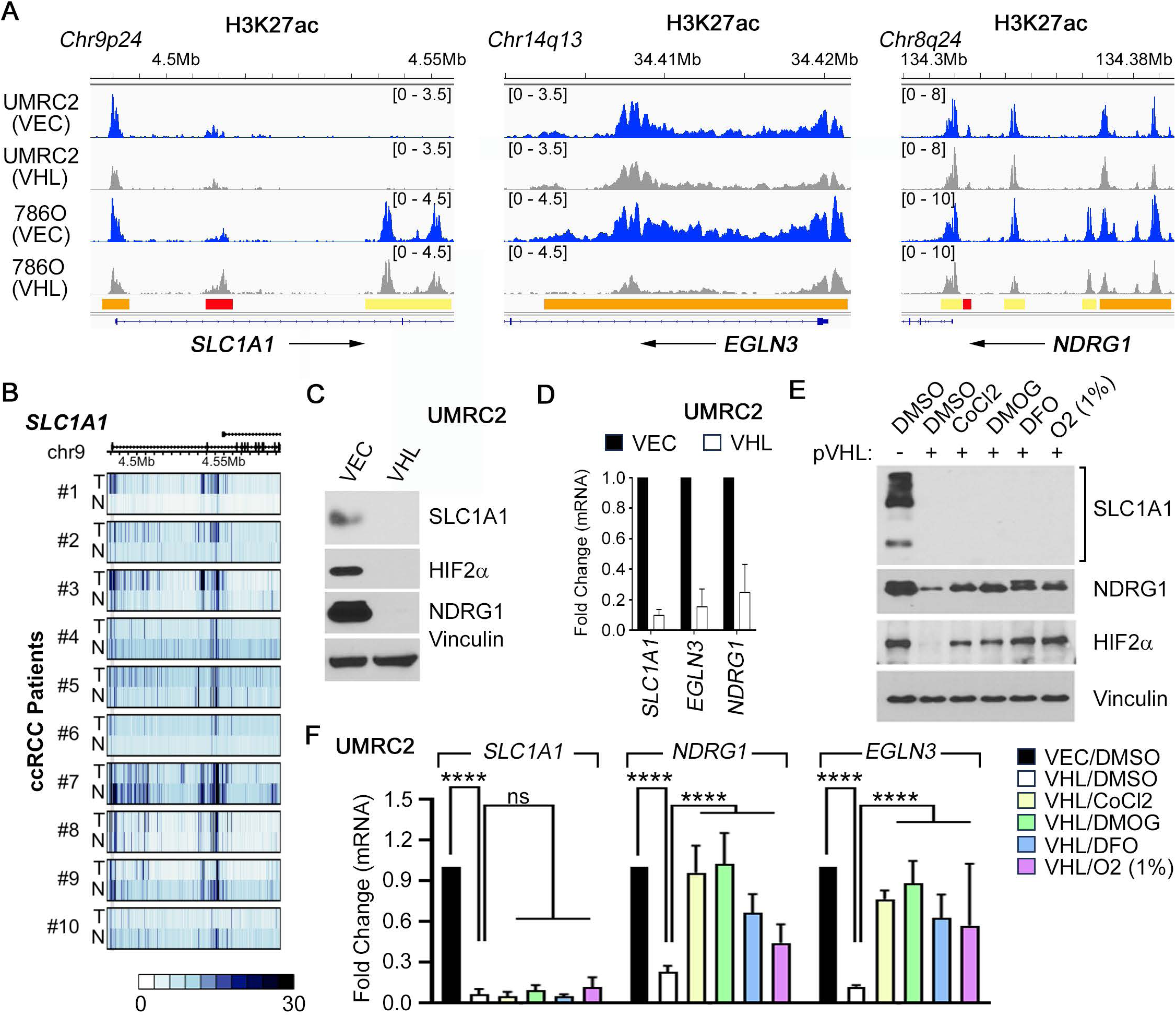
SLC1A1 Expression is pVHL-dependent but HIF-independent. (**A**) H3K27ac signal, plotted using the Integrated Genomic Viewer, at the indicated genomic loci in pVHL-deficient (VEC) and pVHL-proficient (VHL) versions of two ccRCC cell lines (UMRC-2 and 786-O). (**B**) H3K27ac signal at the genomic loci in (A) in either human ccRCC tumors (T) or their paired adjacent normal tissue (N), as indicated. (**C** and **D**) Immunoblotting (**C**) and real-time qPCR (**D**) of the indicated genes in pVHL-deficient (VEC) and pVHL-proficient (VHL) versions of UMRC-2 cells. (**E** and **F**) Immunoblot analysis of the indicated proteins (**E**) and real-time qPCR of the indicated genes (**F**) in pVHL-deficient or pVHL-proficient versions of UMRC-2 cells, as indicated, that were treated for 16 h with the indicated hypoxia mimetic molecules (CoCl_2_, DMOG, or DFO) or environmental hypoxia (1% O_2_).

To address HIF dependence, we subjected pVHL-proficient cells to pharmacological (e.g., DMOG, DFO, and CoCl_2_) or physiological (e.g., 1% oxygen) conditions that attenuate EglN activity. As expected, treatment with hypoxia or these hypoxia mimetics stabilized HIF2α and restored HIF target genes (e.g., *NDRG1* and *EGLN3*) in pVHL-proficient cells. However, these treatments failed to restore SLC1A1 levels (Fig. 4E and F). We then lentivirally infected pVHL-proficient cells to express dual proline to alanine mutant (dPA) versions of either of the two transcriptionally active HIFα isoforms: HIF1α and HIF2α. These mutants are not recognized by pVHL and are thus constitutively active. HIF1αdPA and HIF2αdPA restored the expression of their corresponding target genes (e.g., *DDIT4*, *SLC2A1*/*GLUT1* and *NDRG1*), but failed to restore SLC1A1 expression (Supplementary Fig. S2E and S2F). Finally, using CRISPR/Cas9, we inactivated HIF1β (or ARNT), the obligate binding partner for HIF1α and HIF2α. Unlike the canonical HIF target NDRG1, ARNT loss failed to diminish SLC1A1 expression in pVHL-deficient cells (Supplementary Fig. S2G). Together, these observations demonstrated that the expression of SLC1A1 in ccRCCs was pVHL-dependent, but HIF-independent.

### SLC1A1 Drives Oncogenic Metabolic Reprogramming in ccRCC

Our functional studies suggested that SLC1A1’s oncogenic activity in ccRCC was dependent on its ability to import Asp/Glu (Fig. 3H). Asp/Glu can impact many metabolic pathways. Therefore, we next interrogated the potential metabolic mechanisms underlying SLC1A1’s oncogenicity by measuring the impact of CRISPR/Cas9-mediated *SLC1A1* inactivation using a LC-MS/MS-based targeted metabolomics assay that measures the steady-state abundance of ∼300 polar metabolites representing central carbon, amino acid, and nucleotide metabolism (29). These data were analyzed using the MetaboAnalyst suite (30). As expected, SLC1A1 inactivation decreased the intracellular pools of Asp/Glu, their modified derivatives [e.g., N-acetyl aspartyl glutamate (NAAG), Carbamoyl aspartate (Car-Asp), and Pyroglutamate (PyroGlu)], and their downstream products in the TCA cycle [e.g., α-ketoglutarate (αKG), Malate (Mal), Fumarate (Fum), and Succinate (Suc)]. Interestingly, however, SLC1A1 depletion also reduced the abundance of several metabolites in the nucleotide/1-carbon metabolism pathways (Fig. 5A to E; Supplementary Fig. S3; Supplementary Table S3).

**Figure 5.**
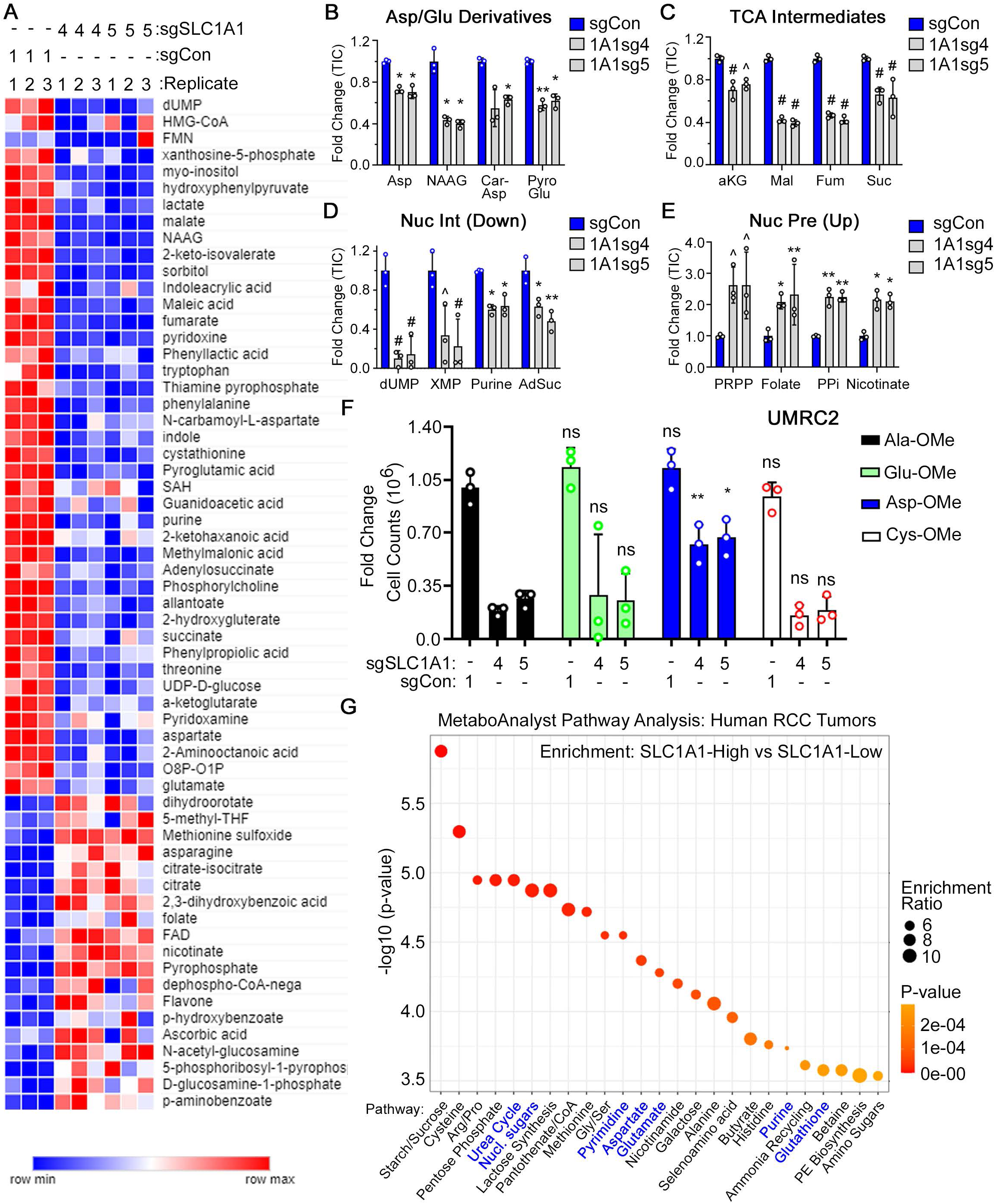
SLC1A1 Function Supports One-Carbon and Nucleotide Metabolism. (**A** to **E**) Heatmap of normalized ion counts (**A**) and fold-change in the abundance of representative metabolites in the indicated pathways (**B** to **E**), as measured using LC-MS/MS analysis from UMRC-6 cells that were lentivirally transduced to express two independent sgRNAs targeting SLC1A1 (1A1sg4 or 1A1sg5) or a non-targeting control (sgCon), as indicated. Metabolites were harvested one day post-selection with puromycin (2 μg/mL) before the emergence of overt cytotoxicity. Ion counts were compared using two-way ANOVA (sgCon vs 1A1sg4 and sgCon vs 1A1sg5) with Dunnett’s multiple comparison test, n=3, *p<0.05; **p<0.01; ^p<0.001; ^#^p<0.0001. (**F**) Normalized cell counts in UMRC-2 cells (relative to sgCon), as measured using the ViCell fourteen days post-selection with puromycin (2 μg/mL), in cells that were supplemented with 100 μM the indicated esterified amino acids beginning three days prior to infection, and then transduced with the indicated sgRNA constructs. Cell counts were compared using two-way ANOVA (all comparisons made to the Ala-OMe arm for the respective genetic perturbation) with Dunnett’s multiple comparison test, n=3, *p<0.05; **p<0.01; ns = non-significant. (**G**) Pathway analysis, done using the Metaboanalyst software suite, to identify metabolic pathways that were over-represented in human ccRCC tumors in the MSKCC patient cohort with high (top-quartile) versus low (bottom-quartile) SLC1A1 mRNA expression.

To address the relative importance of Asp and/or Glu in SLC1A1 function, we performed metabolic rescue experiments. We cultured pVHL-deficient cells in media supplemented with cell-permeable, esterified, versions of SLC1A1’s ligands (e.g., Asp-OMe, Glu-OMe, or Cys-OMe) or a non-specific control (e.g., Ala-OMe). We then lentivirally transduced these cells with CRISPR/Cas9 constructs to inactivate *SLC1A1*. Profound fitness defects were observed upon SLC1A1 loss in cells that were supplemented with cell-permeable versions of Ala, Glu, and Cys. However, cell permeable Asp was sufficient to, at least partially, rescue these fitness defects (Fig. 5F).

We then explored human tumor datasets to validate these cell-based observations. Using steady-state metabolomics and gene expression datasets generated from renal cancer patients at the Memorial Sloan Kettering Cancer Center (MSKCC IRBs 06-107 and 12-237), we first sorted tumors based on *SLC1A1* expression and then compared steady-state metabolite levels in the top quartile (high SLC1A1 expression) to the bottom quartile (low SLC1A1 expression). We found that high-SLC1A1 tumors showed significant enrichment in many of the same pathways that were depleted upon SLC1A1 loss in our cell-based studies, including aspartate, glutamate, and nucleotide metabolism (Fig. 5G). These observations lent physiological credibility to our cell-based metabolic studies and suggested that SLC1A1’s functional importance in ccRCC tumors was dependent on its Asp uptake activity, which likely supports the tumor’s needs for nucleotide metabolism.

### SLC1A1 Represents a Nutrient Carrier Hub that Transcriptionally Governs Solute Import

What might be the functional relevance of high *SLC1A1* expression in human tumors? To begin addressing this question, we annotated TCGA renal tumors based on *SLC1A1* gene expression and, as a control, expression of the canonical HIF target gene, *CA9*. We noted that, whereas CA9 expression was predominantly limited to ccRCCs, SLC1A1 expression was more widespread (Supplementary Fig. S4A and S4B). We then focused on the ccRCC subset and performed Gene Set Enrichment Analysis [(GSEA) (31)] on tumors annotated into the top versus bottom quartile of *SLC1A1* or *CA9*. High *CA9* expression was associated with signatures of HIF/Hypoxic activity (Fig. 6A to C; Supplementary Table S4); however, consistent with the HIF-independent expression of *SLC1A1*, we found *SLC1A1* expression strongly correlated with widespread metabolic reprogramming but not with Hypoxic programs (Fig. 6D to F; Supplementary Table S5). Analyzing the top 100 upregulated genes associated with high *SLC1A1* in both the TCGA and the MSKCC cohorts using functional interaction tools, such as StringDB, we found that elevated expression of *SLC1A1* was associated with higher expression of many other solute carrier proteins (e.g., ∼25 of the top 100 differentially expressed genes were SLCs) (Fig. 6G; Supplementary Table. S6; Supplementary Fig. S4C). Finally, we noted that restoration of *SLC1A1* in pVHL-proficient cells, which was sufficient to promote tumorigenesis *in vivo*, did not restore expression of HIF target genes, such as *EGLN3* and *NDRG1* (Fig. 6H). Therefore, *SLC1A1* represents a dysregulated – HIF-independent – metabolic hub in ccRCC.

**Figure 6.**
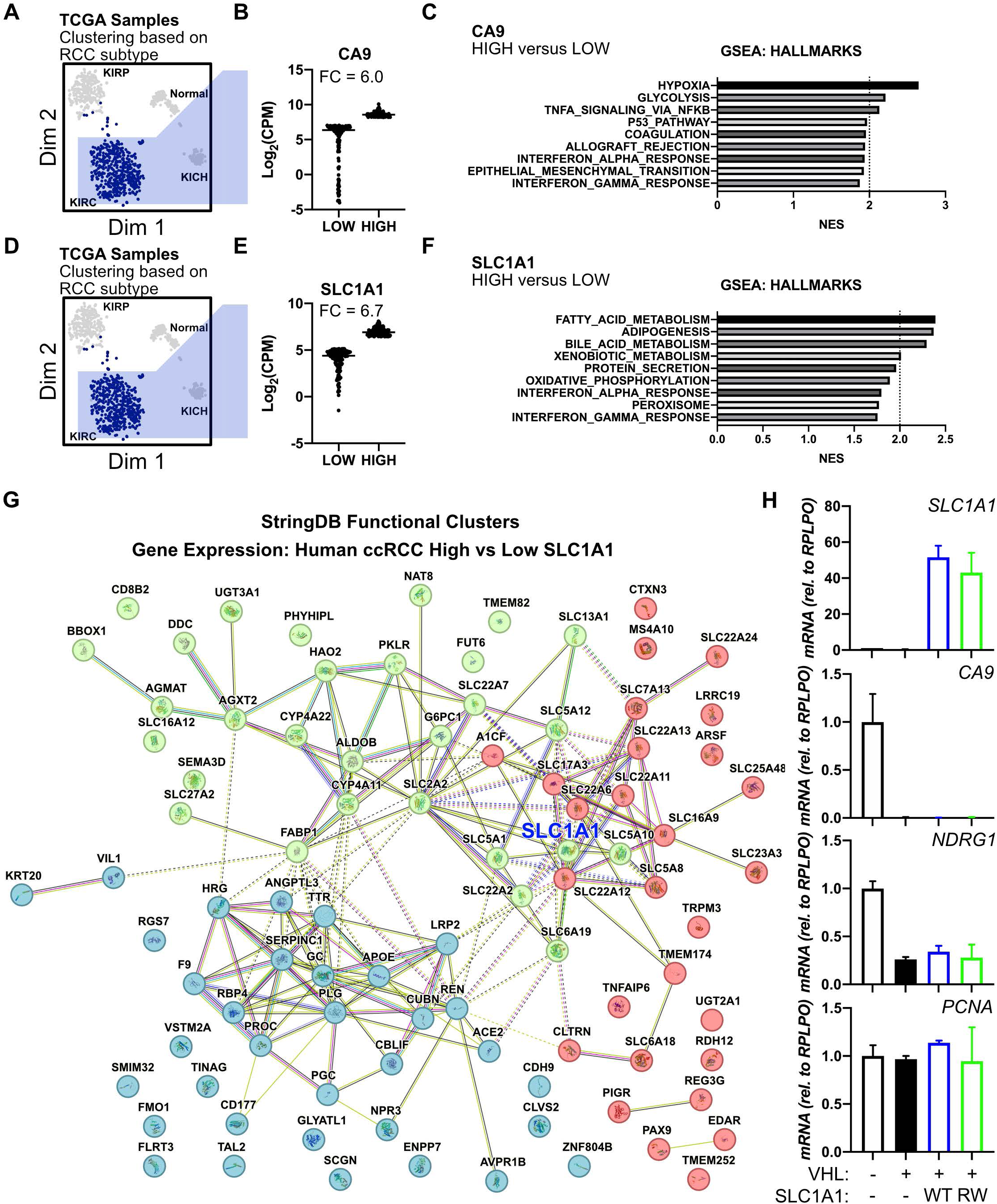
SLC1A1 Represents a Dysregulated, HIF-independent, Metabolic Hub in ccRCC. (**A** to **F**) Principal components analysis and T-Stochastic Neighbor Embedding, fold change in mRNA abundance, and Geneset enrichment analysis, using TCGA human renal cancer data (n=135), for *CA9* (**A** to **C**) or *SLC1A1* (**D** to **F**), respectively, as indicated. (**G**) Functional interaction analysis, using StringDB, for the top 100 genes that are upregulated in tumors with high *SLC1A1* expression. (**H**) Realtime-qPCR analysis of the indicated genes using UMRC-2 cells that were first transduced to express pVHL or VEC (control), selected in Blasticidin (10 µg/mL) for 7 days, then transduced to express either wild-type SLC1A1 (WT), R445W mutant SLC1A1 (RW), or VEC control (-). Cells were selected with Puromycin (2 µg/mL) for 3-5 days before sample collection.

### SLC1A1 Activity Confers Actionable Metabolic Vulnerabilities in ccRCC

Our genetic studies suggested that pharmacological inhibition of SLC1A1 could be a viable strategy to target ccRCCs. In preliminary studies, we confirmed that SLC1A1 was most prominently expressed only in the brain and kidney tissue (Supplementary Fig. S5A). Additionally, literature searches confirmed that homozygous *Slc1a1* knockouts in mice lack any significant growth defects under normal conditions (32). Finally, humans with loss-of-function germline mutations in SLC1A1 show limited clinical symptoms except for the urinary excretion of Asp/Glu, which is consistent with SLC1A1’s role in amino acid reabsorption in the kidney (27). This tissue-restricted expression and lack of phenotype in normal tissue supports the existence of a therapeutic index when using SLC1A1 inhibitors.

SLC1A1 is of significant interest in neuroscience because it modulates glutamatergic neuronal transmission by clearing post-synaptic extracellular glutamate (33, 34). Previous medicinal chemistry campaigns have led to SLC1A1 inhibitors based off two scaffolds. The first scaffold class maintains an aspartate-like backbone (35–39), which lacks specificity and pharmaceutical properties for systemic dosing. The second scaffold is derived from bicyclic imidazopyridine anilines (40), which maintains a steep Structure-Activity Relationship (SAR) profile requiring a furan ring for activity against SLC1A1, as represented by compound <u>3e</u> (Cmpd <u>3e</u>) (Supplementary Fig. S5B). This molecule not only suffers from poor solubility, but the presence of the electron-rich heterocyclic furan also makes it susceptible to rapid clearance by the cytochrome P450 (CYP) enzymes and limits Cmpd <u>3e</u>’s *in vivo* utility.

Despite these limitations, we experimentally characterized the utility of Cmpd <u>3e</u> in ccRCC, hoping that this compound could be a useful non-amino acid tool compound that may lead to a scaffold for eventual medicinal chemistry. We synthesized Cmpd <u>3e</u> using the reported Groebke−Blackburn−Bienaymé condensation. After quality control steps to confirm desired purity and structure (Supplementary Fig. S5C), we tested the direct binding of Cmpd <u>3e</u> with SLC1A1 using a Cellular Thermal Shift Assay (CETSA). We treated membrane-enriched fractions harvested from renal cells expressing V5-tagged SLC1A1 with a relatively high-dose of Cmpd <u>3e</u> (100 µM) or DMSO (as control) and then measured SLC1A1 abundance at different temperatures. These studies showed that exposure to Cmpd <u>3e</u> increased SLC1A1 stability, in line with direct interaction of this molecule (Supplementary Fig. S5D). We then tested the effects of Cmpd <u>3e</u> on RCC cells that were sensitive to acute SLC1A1 inhibition (e.g., A498), as observed using CRISPR/Cas9-dependent gene editing. Consistent with our genetic experiments, we noted significant cytotoxicity in A498 cells (IC_50_: ∼0.9 µM). In contrast, UMRC2 cells, which showed a delayed response to genetic inactivation of SLC1A1, were largely insensitive to Cmpd <u>3e</u> at early time points (IC_50_: ∼30 µM) (Supplementary Figure S5E and S5F). The Glutaminase inhibitor CB-839 was studied in parallel as a positive control based on prior reports (41), and showed comparable cytotoxic effects in both cell lines.

We then evaluated the disposition and Pharmacokinetic (PK) properties of Cmpd <u>3e</u> *in vivo* after an acute systemic intraperitoneal (IP) administration of Cmpd <u>3e</u> in NSG mice at 10 mg/kg (Supplementary Fig. S5G). We found Cmpd <u>3e</u> has a plasma C_max_ of only ∼90 nM and a relatively short half-life (∼2.5 hrs) based on its plasma drug concentration time course (Supplementary Fig. S5H). This observation *in vivo* is consistent with *in vitro* microsomal metabolic stability profiles in human and rat, which indicate high turnover (human and rat intrinsic clearance > 80% hepatic, human CL_int_ > 142 mL/min/kg, rat CL_int_ 495 mL/min/kg) (Supplementary Fig. S5I). Interestingly, Cmpd <u>3e</u> does accumulate in the kidney to some degree (C_max_: 1.25 µM) but preferentially accumulates in the CNS (C_max_: 5.84 µM) (Supplementary Fig. S5H). In summary, Cmpd <u>3e</u> displayed moderate potency but less than ideal properties for systemic dosing where oral delivery would be preferred, thus severely limiting its utility in tumor efficacy studies.

### SLC1A1 Blockade Offers Clinically relevant Therapeutic Opportunities in ccRCC

Given the challenges with using Cmpd 3e *in vivo*, we addressed the clinical relevance of SLC1A1 using genetic tools both *in vitro* and *in vivo*. HIF activation, both upon pVHL loss and in physiological hypoxia, rewires the TCA cycle and drives Gln in the (reverse) reductive direction to generate Citrate (42, 43). This reductive utilization of Gln is driven by Glutaminase, which converts Gln into Glu. Glutaminase blockade (e.g., using CB-839) thus not only diminishes Glu pools but also decreases Asp produced from TCA intermediates such as oxaloacetate and represents a metabolic vulnerability in kidney cancer (Supplementary Fig. S5F) (41, 44).

We reasoned that Asp/Glu uptake by SLC1A1 would impact Glutaminase dependency in ccRCC (Supplementary Fig. S6A). Specifically, we hypothesized that SLC1A1 overexpression (and consequent elevated Asp/Glu pools) would reduce dependence on Glutaminase, whereas inactivation of SLC1A1 (and consequent depleted Asp/Glu pools) would sensitize cells to Glutaminase blockade. Indeed, we observed that overexpression of wild-type SLC1A1, but not the functionally-dead RW mutant, conferred resistance to CB-839 (Supplementary Fig. S6B and S6C). Conversely, CRISPR/Cas9-mediated inactivation of SLC1A1 sensitized ccRCC cells to CB-839 treatment (Supplementary Fig. S6D and S6E).

Diminished Asp upon Glutaminase inhibition leads to a depletion of intracellular nucleotide pools and causes elevated replicative stress and DNA damage in pVHL-deficient cells (44). Therefore, we reasoned that, restoration of cellular Asp/Glu via SLC1A1 would counteract these effects. Indeed, using nuclear γH2AX foci as a marker of DNA damage, we found markedly lower DNA damage upon CB-839 treatment in ccRCC cells expressing WT SLC1A1 versus those expressing the functionally-dead SLC1A1 RW mutant (Supplementary Fig. S6F to S6H).

Lastly, we addressed the importance of SLC1A1 dependency *in vivo* using genetic tools. We engineered constructs that expressed sgRNAs targeting SLC1A1 (or a non-targeting control) under the control of doxycycline (Dox), using the TLCv2 lentiviral backbone (Supplementary Fig. S6I). UMRC-2 cells, transduced with these constructs, were expanded and inoculated in sub-cutaneous xenografts in NSG mice. As expected, tumor formation for the first several weeks in the absence of Dox did not differ between the two experimental cohorts (sgCon versus sgSLC1A1) (Supplementary Fig. S6J).

Once tumors reached ∼100 mm^3^ all the mice were switched to Dox-containing chow for a period of fourteen days. Interestingly, despite the relatively weak knockdown efficiency with this vector, we noted that SLC1A1 inactivation measurably decreased tumor growth, establishing the importance of SLC1A1 as an oncogenic target in ccRCC (Supplementary Fig. S6J and S6K).

### Elevated SLC1A1 Expression Correlates with Metastatic ccRCC

Finally, we addressed the clinical relevance of SLC1A1 expression as a biomarker in human renal tumors. We first compared the expression of SLC1A1 in some human ccRCC tumors (3 female versus 4 male patients; Supplementary Table S7) and their paired normal adjacent tissue, banked upon consent from patients at Cleveland Clinic (IRB 4639). Surprisingly, we found that bulk expression level of SLC1A1 was lower in tumors compared to normal tissue (Supplementary Fig. S7A). This analysis was sub-optimal because of: (**1**) the small sample numbers; (**2**) the technical differences in extracting membrane proteins from normal versus stromal-rich tumor tissue; (**3**) the likely presence of non-tumor cells in bulk tumors; and (**4**) the presence of necrotic, non-viable, tumor regions. Therefore, we developed IHC assays to reliably detect SLC1A1 *in situ* and enable a comparison of SLC1A1 expression in normal versus tumor tissue. Using cell lines that express either low SLC1A1 (UMRC-2; pVHL) or high SLC1A1 (UMRC-2; VEC), we optimized a SLC1A1 staining protocol in FFPE samples (Supplementary Fig. S7B and S7C).

In normal human kidney tissue, stained using this optimized IHC protocol, we did not observe any SLC1A1 staining in the hypoxic renal medulla. But notable SLC1A1 expression was seen on the apical, lumen-facing, side of the renal cortical epithelium, consistent with SLC1A1’s role in amino acid reabsorption. In tumor tissue, however, SLC1A1 expression was redistributed throughout the entire cell surface (Supplementary Fig. S7D). This difference in localization likely marks the loss of apico-basal polarity in renal tumors, but also perhaps an altered functional role for SLC1A1 in amino acid uptake from the ccRCC tumor microenvironment.

We then analyzed a large tumor/normal tissue microarray (TMA) panel generated from renal cancer patients seen at DFCI and BWH (IRB 01-130). We qualitatively scored for membranous SLC1A1 staining intensity (+1 to +3) and the proportion of tumor stained. The product of these scores was cumulatively reported as an H-Score, a standard approach that delivers scores between 100 to 300, which is employed to measure staining intensity by pathologists. We noted that benign renal oncocytomas and renal chromophobe tumors had no detectable membranous staining of SLC1A1 (H-score = 0, Fig.7A and B). In contrast, SLC1A1 expression was detected, albeit at low levels, in both clear cell RCC and papillary RCC tumors (H-score = ∼120-150, Fig.7A and B). These expression patterns were also consistent with TCGA data, which indicated significantly higher *SLC1A1* mRNA expression in both ccRCC and pRCC, compared to the chromophobe tumors (Fig.7C; Supplementary Table S8).

**Figure 7.**
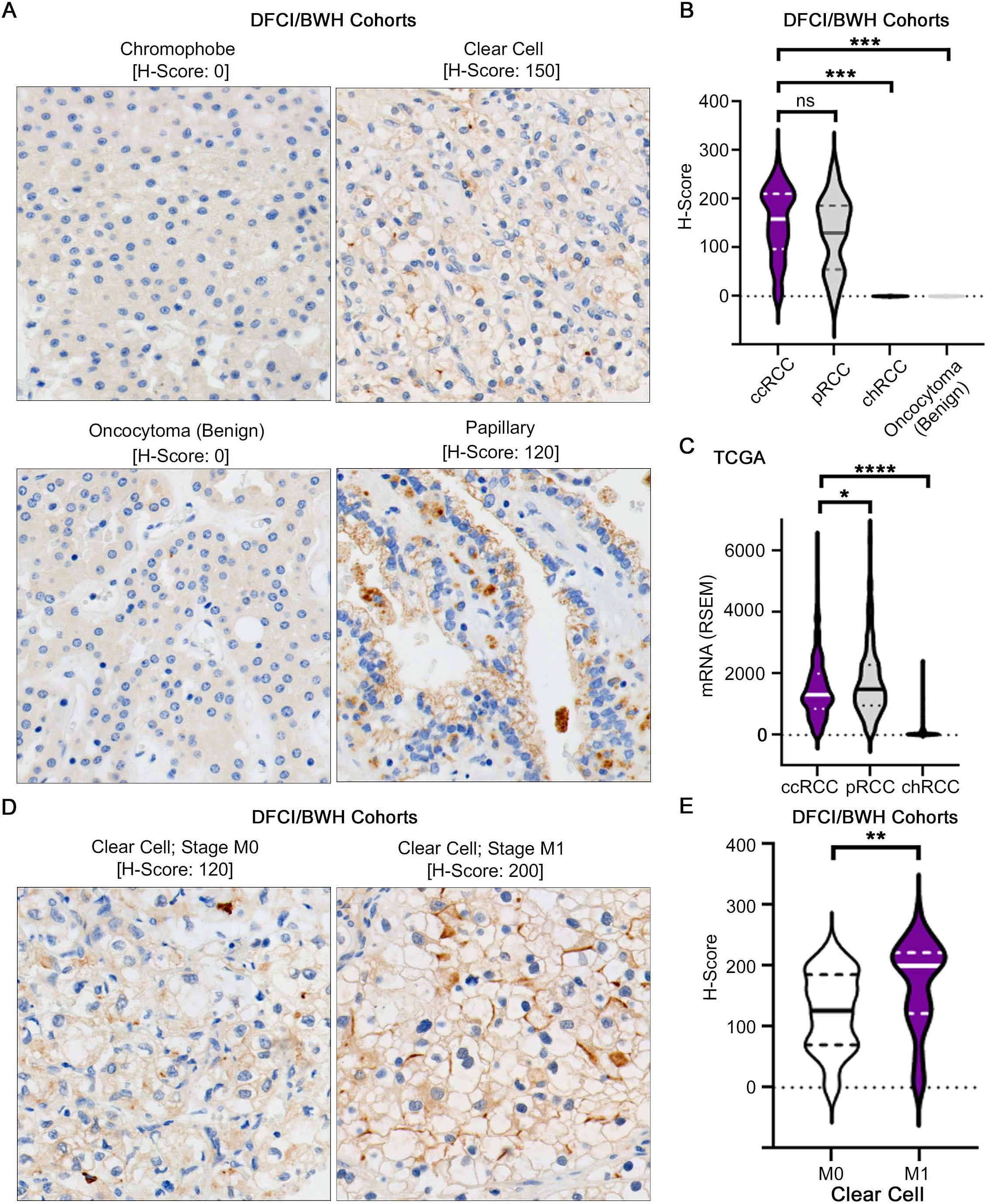
Human Renal Tumor Analysis Identifies Clinical Correlates of SLC1A1. (**A** and **B**) Representative photomicrographs (**A**) and pathological quantification of the H-score representing SLC1A1’s membrane expression (**B**) in sections of the indicated human renal tumors. In (**B**), H-scores in all other cohorts were compared to the ccRCC arm using two-way ANOVA with the Dunn’s multiple comparison test; ccRCC (n=82); pRCC (n=13); chRCC (n=5); Benign tumors (n=7). ***p<0.001; ns = non-significant. (**C**) Comparison of SLC1A1 mRNA expression in the indicated renal cancer subtypes (ccRCC n=510; pRCC n=283; chRCC n=65) with data mined from the TCGA, using the two-way ANOVA with Dunn’s multiple comparison test; *p<0.05; ****p<0.0001. (**D** and **E**) Representative photomicrographs (**D**) and pathological quantification of the H-score representing SLC1A1’s membrane expression (**E**) in sections of ccRCC tumors that represented either metastatic (M1) or non-metastatic (M0) disease states. H-scores in the M0 (n=34) and M1 (n=48) cohorts were compared using the Mann-Whitney test; **p<0.01.

Importantly, when sorted by stage, higher SLC1A1 expression associated with advanced stage 4 disease (H-score = ∼200, Supplementary Fig. S7E). Consistent with this observation, we found discernibly higher SLC1A1 expression in metastatic (M1) compared to non-metastatic (M0) ccRCCs (Fig.7D and E). In summary, these studies on clinical material established an unexpected redistribution of SLC1A1, away from the apical surface, on the membranes of tumor cells and an association of SLC1A1 expression with metastatic disease.

## DISCUSSION

Epigenetic dysregulation often drives widespread transcriptional changes, which poses technical challenges in the identification of downstream oncogenic targets. A typical strategy to fish for such oncogenic targets is deploying a negative-selection or ‘dropout’ genetic screen, which scores for the loss of sg/shRNA representation in unhealthy/dying cells *in vitro*. This approach tends to be plagued with false-positive scoring, driven by the off-target cytotoxicity of these genetic tools. Performing this analysis *in vitro* also limits the discovery of physiologically relevant metabolic targets, whose dependency could be profoundly influenced by the nutrient availability in the tumor environment.

To mitigate these concerns, we made several modifications to our target discovery pipeline. First, instead of a ‘dropout’, we designed a positive-selection approach, which scored a gene’s ability to confer tumorigenic potential. Second, we performed our screen *in vivo*, to enable the identification of physiologically relevant metabolic targets. Third, we rigorously validated the hits from our initial screen using secondary *in vivo* assays and cell-based studies. All primary observations relied on multiple ccRCC cell lines, used multiple sgRNAs targeting SLC1A1, were confirmed to be on-target using rescue experiments, and were supported by validations in human tumor samples. In summary, this analytical pipeline represented a rigorous experimental strategy that led to the discovery of an epigenetically dysregulated metabolic target.

The role of SLC1A1 is poorly understood in cancer biology. Much of the biological characterization of SLC1A1 has been done in neurological models, where this transporter is known to limit neurotransmission by removing Glu from post-synaptic space. However, some evidence for SLC1A1’s oncogenic role has been recently reported in human tumors, including lung (45), T-cell lymphomas (46), and gliomas (47). The importance of Asp/Glu uptake in tumor biology is also supported by the oncogenic importance of other SLC1A1 paralogs (e.g., SLC1A3) in hypoxic tumors (48). Consistent with this growing body of work, our studies highlight how SLC1A1 promotes ccRCC carcinogenesis and justify the utility of SLC1A1 blockers as anti-cancer agents.

Metabolic studies in human renal tumors have shown the significant elevation of 1 carbon intermediates (versus normal adjacent tissue), which is speculated to support nucleotide biosynthesis in these tumors (49). Our studies suggest that SLC1A1’s functional role as an Asp/Glu importer provides the key carbon skeleton for the biosynthesis of nucleotide intermediates. Additionally, our studies also establish that SLC1A1 function could impact the cellular response to other metabolic interventions, in particular interventions aimed at the Gln utilization pathways.

The functional role of pVHL as an E3 ligase that targets HIF2α for proteolysis has defined almost all our drug-discovery efforts in kidney cancer, including the discovery of TKIs that target the HIF-target gene VEGF; targeting HIF-dependent nutrient-sensing pathways, such as mTOR; and most recently the HIF2α inhibitors. Despite its well-recognized role as a driver in ccRCC, HIF2 blockade is not sufficient to treat all sporadic ccRCCs, suggesting the presence of genetic heterogeneity and/or the existence of pVHL-dependent, but HIF-independent, oncogenic mechanisms. Our discovery shows that the expression of metabolic regulators, such as SLC1A1, represents one such HIF-independent pathway. It will be important in the future to understand the relative importance of SLC1A1 as a target in both HIF2α-dependent and independent ccRCC tumors as well as the mechanism by which pVHL regulates SLC1A1. Importantly, our findings here show that SLC1A1 function was relevant in both HIF2α-dependent cells (e.g., A498) and HIF2α-independent cells (e.g., UMRC-2).

Our analysis of bulk tumors suggested, surprisingly, that SLC1A1 expression was lower in tumors compared to normal adjacent tissue. However, this analysis had some biological and technical caveats. Differences in SLC1A1’s localization in normal versus tumor tissue could impact the extraction efficiency of this membrane-embedded protein, especially in stromal-rich tumors. Additionally, the presence of some non-tumor cells, which we also found expressed SLC1A1 on their apical surface, is a potential confounder in measurements from bulk tumors. Yet, interpreting this anomalous finding at face value, we speculate that the loss of cellular polarity redistributes and decreases SLC1A1 protein abundance in renal tumors, thereby converting them into quasi-auxotrophs with increased dependence on the residual activity of SLC1A1. Targeting this residual SLC1A1 thus proves deleterious in ccRCCs.

One caveat of our study was that the existing SLC1A1 inhibitor lacked the pharmacological properties to effectively target the transporter *in vivo*. We did, however, note cytotoxic responses in cell-based studies, where pharmacological SLC1A1 blockade phenocopied *SLC1A1*’s genetic inactivation. Future studies could focus on the development of Cmpd <u>3e</u> analogs that improve its metabolic stability, offer oral delivery, and are non-brain penetrant to avoid targeting SLC1A1 in the CNS. Altogether, our studies establish a strong rationale to launch future medicinal chemistry campaigns that develop novel SLC1A1 blockers.

## METHODS

### Cell Lines

A-498 (RRID: CVCL_1056) and 786-O (RRID:CVCL_1051) renal carcinoma cells were obtained from American Type Culture collection. UMRC-2 (RRID: CVCL_2739) and UMRC-6 (RRID: CVCL_2741) cells were obtained from Dr. Bert Zbar and Dr. Marston Linehan (National Cancer Institute). HEK293T (RRID:CVCL_0063) was obtained from the Lerner Research Institute’s Cell Media Core. 293FT cells were procured from Thermofisher (ThermoFisher, R70007, RRID:CVCL_6911). All cells were maintained in Dulbecco’s Modified Eagles Medium (DMEM) (Life Technologies 11995073) supplemented with 10% Fetal Bovine Serum (FBS) (Life Technologies 10437-028), and 1X Penicillin-Streptomycin (Life Technologies 15140163). Cells stably expressing cDNAs or sgRNA were generated by lentiviral infection followed by selection with Puromycin (2 µg/mL) or Blasticidin (10 µg/mL), as appropriate. All cells were grown at 37°C in 5% CO_2_.

### Plasmids

The ORF library encoding the candidate oncogenes and miscellaneous controls was obtained from the Broad Institute’s Gene Perturbation Platform in the pLX317 backbone, which has a unique 20 nt barcode associated with each gene. The barcode descriptions are provided in Supplementary Table S2. The CRISPR/Cas9 constructs to target *SLC1A1* were based on sequences provided in the Gecko library and generated by annealing the appropriate sgRNAs (described in the Oligo list) and cloning them into BsmBI-digested pLentiCRISPRv2 plasmids (RRID: Addgene_52961).

### Lentiviral Infections

Lentiviruses were generated by packaging desired constructs into either 293FT or 293T packaging cells. Typically, the packaging cells were seeded into 6 cm plates (∼1.8E06 – 2.6E06 cells), attached overnight, and then transfected. The standard transfection mixture included 3 μg of total DNA [1.5 µg lentiviral plasmid + 1.5 µg helper plasmids - psPAX2 (RRID: Addgene_12260) and pMD2.G (RRID: Addgene_12259) mixed in a 3:1 ratio], combined with 9 µl lipofectamine 2000 (Invitrogen 11668019). Transfection mixtures were incubated at room temperature for 15 mins, added dropwise onto cells, and then incubated for sixteen hours. Following this, media was changed and viral supernatant was collected at 48- and 72-hours post transfection and combined.

Lentiviral supernatants were filtered (0.45 μm), aliquoted, and then frozen at -80°C. Cells were seeded for viral infections into 6-well plates (150,000 – 200,000 per well). The following day, cells were transduced with 0.5 mL of the lentiviral supernatant in the presence of either 8 μg/mL polybrene (A-498 and 786-O cells) or 2 μg/mL polybrene (UMRC-2, and UMRC-6 cells). The plates were centrifuged at 300× *g* for 40 min and returned to the incubator for 8 hours after which media was replaced and cells were grown for an additional 24 hours prior to selection with the appropriate antibiotic.

### Generation of *SLC1A1* Mutant Constructs

The SLC1A1 gene was sub-cloned from the pLX317-SLC1A1 construct obtained from the Broad Institute (Clone#) using Gateway cloning (Invitrogen) into the pDONR223 backbone to generate the pEntry-SLC1A1 construct. Mutations encoding I395Δ, R445W, and mismatches that conferred resistance to sg5 were generated by site-directed mutagenesis (Quickchange II XL; Agilent 200521) using the oligos described in the primer list and verified by sanger sequencing. Expression constructs were generated by cloning the SLC1A1 mutant constructs into the pLX304 destination vector (RRID: Addgene_25890) using Gateway cloning (Invitrogen). The plasmids were packaged into lentiviral particles, as described above. For rescue experiments, cells were first engineered to express sgRNA-resistant versions of either WT or mutant SLC1A1 (or GFP, as a control), selected with blasticidin (10 μg/mL, 7-10 days), and then lentivirally transduced with sgRNAs targeting either SLC1A1 or a non-targeting control and selected with puromycin (2 µg/mL, 3 days).

### Primer List

#### Mutagenesis Primers

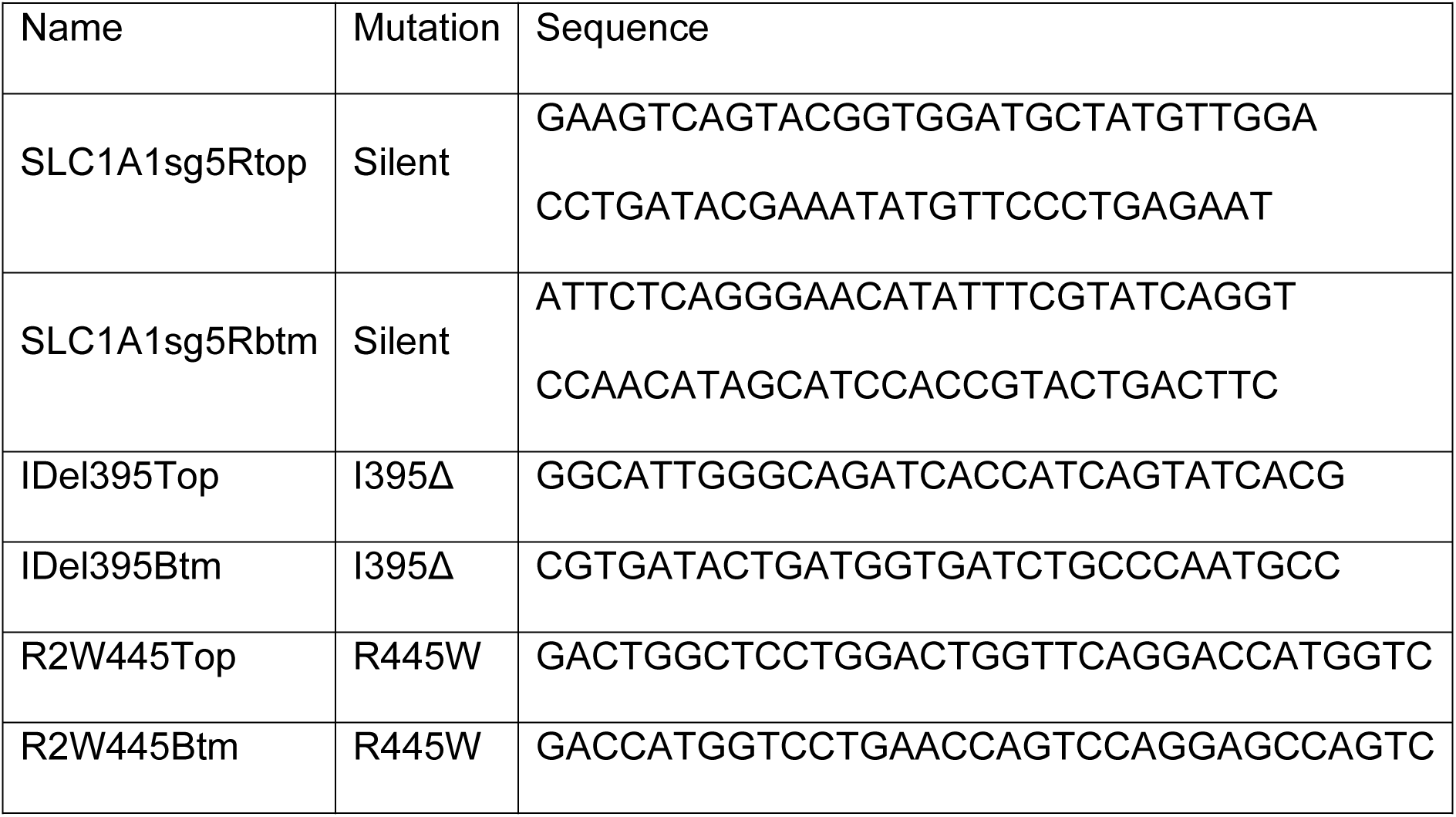

#### sgRNA sequences

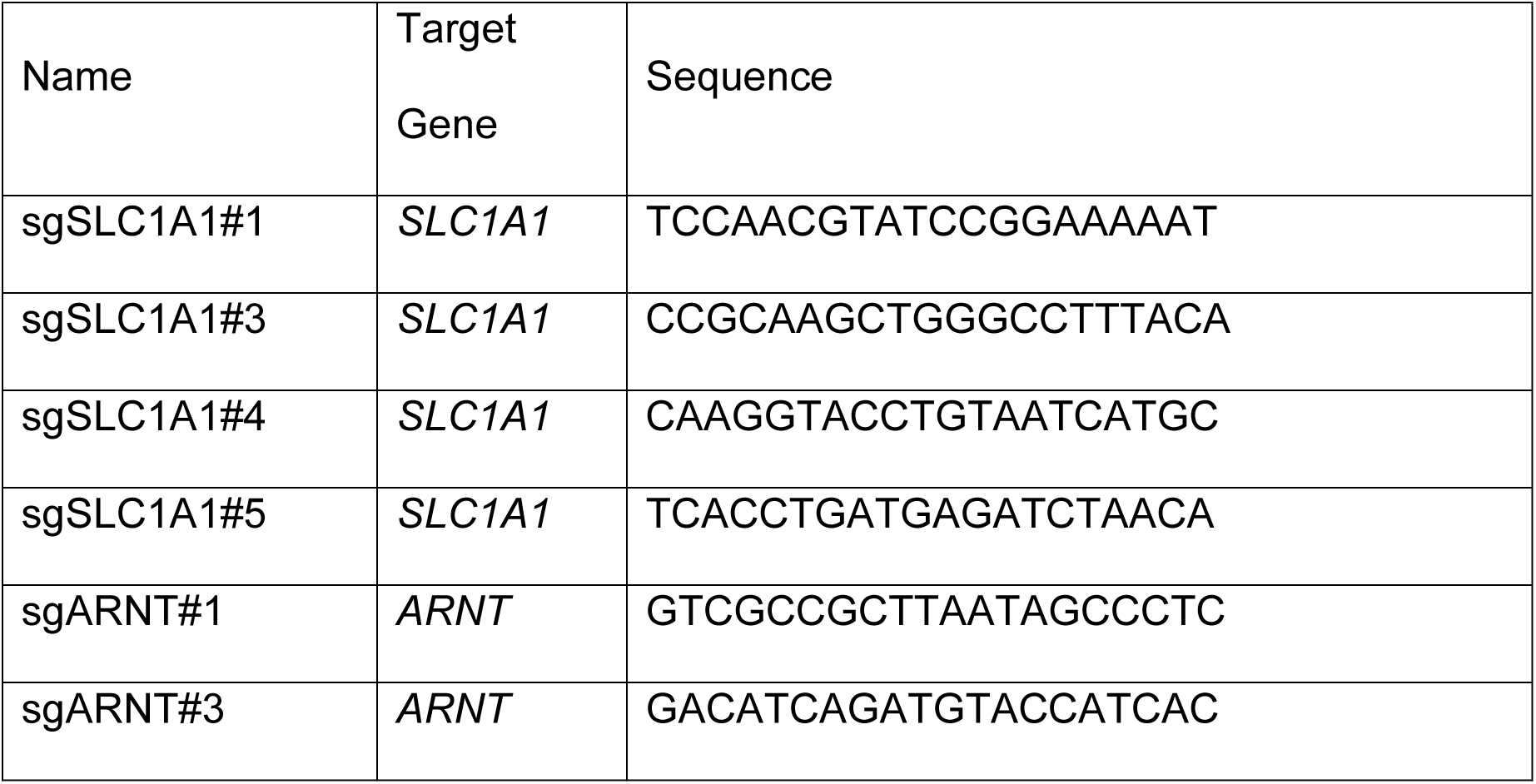

#### qPCR Primer sequences

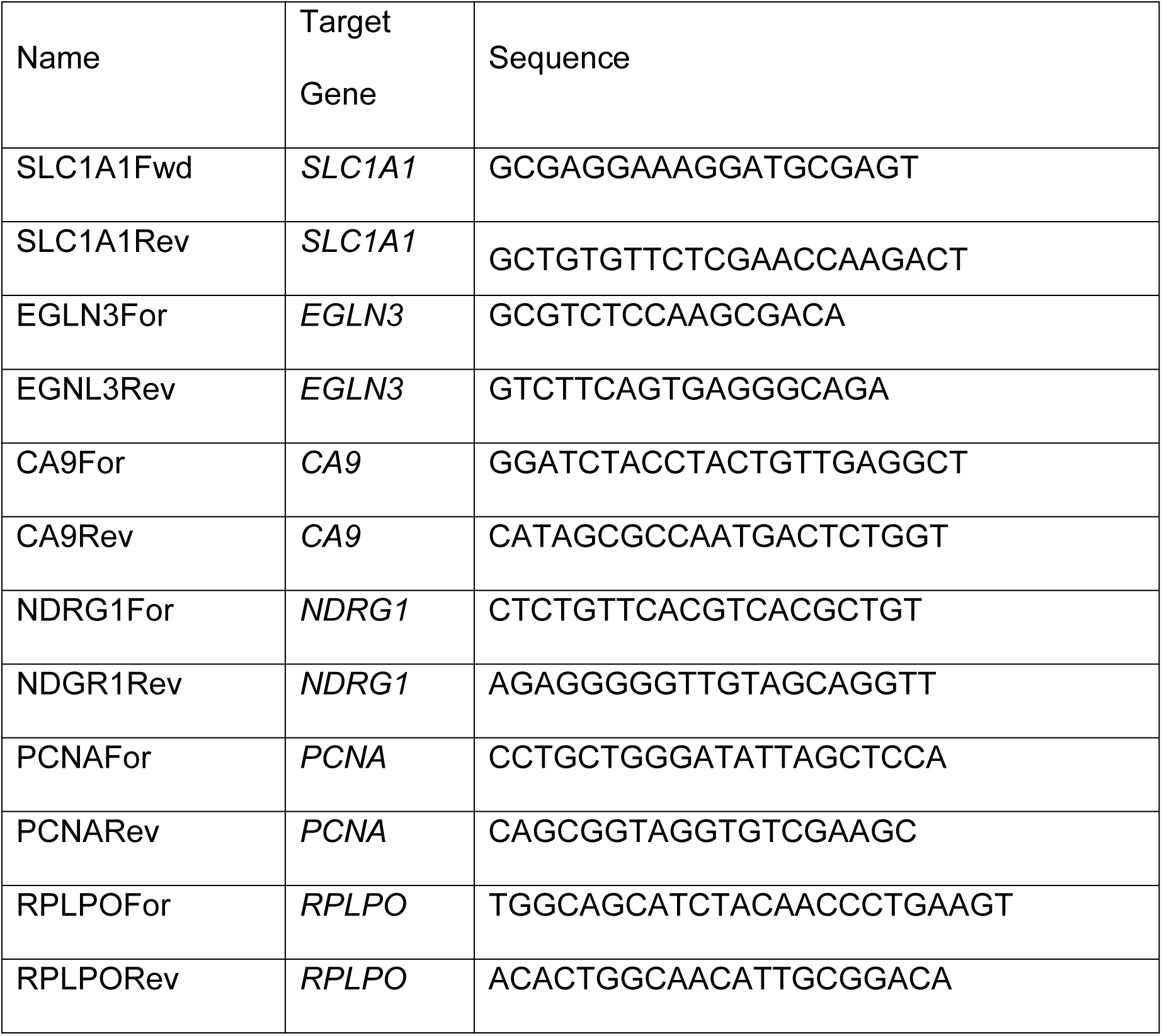

### ChIP-Seq Analysis

Isogenic human ccRCC lines, lentivirally transduced to express HA-pVHL (VHL) or Empty Vector (VEC) control, were expanded. ∼20 million cells (typically 3 x 15 cm dishes) were washed twice with ice-cold 1X PBS and then fixed using 1% methanol-free formaldehyde (Thermofisher 5701) in 1X PBS. Excess formaldehyde was quenched by addition of 125 mM Glycine. Fixed cells were harvested and lysed in SDS lysis buffer [1% SDS, 10mM EDTA, 50mM Tris-HCl (pH 8) + protease inhibitor cocktail (PIC, cOmplete Mini, Roche)] and sonicated under optimized conditions that yielded an average DNA length of ∼300 bp, typically 6 cycles of 30 secs each using a microprobe sonicator at 30% output. Chromatin was quantified and 10 μg total chromatin was used for each H3K27ac ChIP. ChIP-Seq libraries were prepared using the Illumina TruSeq library kit and analyzed using two independent strategies. In one approach, we used the analytical pipelines described previously (50). In another, significant peaks were called using MACS2 broadpeaks (Verson 2.2.7.1) and superenhancers were identified using Homer findpeaks (Version 3.12) (-style super -minDist 12500). Bam files were normalized for sequencing depth (readCount) and log_2_ fold changes between VHL and VEC samples were determined using Deeptools bamCompare (Version 3.5.1). Finally, log_2_ fold changes at significant peaks and superenhancers were determined using Encode (Version 130.2) bigWigAverageOverBed. Data from both approaches largely overlapped and is provided in GSE241864.

For patient samples, raw read files of H3K27ac ChIP-seq, published previously (23), were downloaded from NCBI SRA. Reads with low quality were trimmed using Cutadapt (Version 4.0) and quality cutoff >20. Reads smaller than 20 bp were removed. Trimmed reads were aligned to hg19 using RNA STAR (Version 2.7.8a) and only uniquely mapped reads with mapQ ≥ 60 were used for downstream analysis. Multiple bam files representing the same sample were merged using Samtools merge (Version 1.16.1). Merged bam files were normalized for sequencing depth (SES) and background subtracted using Deeptools bamCompare (Version 3.5.1) against input samples. Log_2_ fold changes between tumor and normal samples were then calculated using Deeptools bigwigCompare. Karyotype plots showing the abundance of H3K27ac at specific genomic regions of interest were generated using KaryoplotR.

### RNA-Seq Analysis

Total RNA, extracted from the isogenic (pVHL-proficient versus deficient) ccRCC cell lines using Trizol reagent (Life Technologies), was used for library preparation with or without polyA enrichment. Following standard QC (e.g., quantification on Qubit and RIN determination on a Bioamalyzer), cDNA libraries were prepared using Illumina TruSeq RNA Library Prep Kit with 50-100 ng of total RNA and sequenced on the Illumina HiSeq 4000. Raw sequences (.fastq files) were quality control tested using FastQC (FastQC, RRID:SCR_014583). The 40 bp single end reads had on average high-quality metrics (>30 Phred score) and nucleotide distributions. Reads with Phred (Phred, RRID:SCR_001017) scores <28 were trimmed using Cutadapt (cutadapt, RRID:SCR_011841) v1.16. While FPKM values derived using Illumina’s BaseSpace RNA-Seq Alignment workflow were initially used to determine log_2_ fold-changes, the RNA-Seq datasets were also reanalyzed using more contemporary pipelines. To this end, reads were aligned to the UCSC hg19 build of the human transcriptome using HISAT2 (HISAT2, RRID:SCR_015530) v2.1.0 and total read counts for genomic features (i.e., genes) were measured using featureCounts (featureCounts, RRID:SCR_012919) v1.6.4. Counts were filtered, normalized (Trimmed Mean of M-values), and differential expression was determined using EdgeR (v3.12).

### ORF Library Screen and Validation

pVHL-proficient UMRC-2 cells were transduced in four independent infections (R1 to R4) with the ORF library (described in Supplemental Table S2) and selected using Puro (2 µg/mL). Cells were harvested and approximately 10% were set aside to measure initial barcode representation (T0), whereas the remaining 90% of the cells were expanded and injected subcutaneously into mice on both flanks at 5E6 cells per injection into immunodeficient NCR^nu/nu^ mice giving two tumor injections per biological replicate (totaling eight injections). Following 8-12 weeks of growth, tumors were harvested, cut into two pieces to either be fixed in 10% formalin or processed for downstream sequencing (Tend). The barcode region of the integrated lentiviral cassettes was amplified from the genomic DNA of T0 and Tend samples, indexed, and sequenced on a Illumina Miseq sequencer. Read counts from each sample was normalized to the total counts from the corresponding sample, and then representation in Tend samples was compared to the corresponding T0 to identify candidate genes that showed enrichment (log2fold change > 1.0) in at least two of the five tumors. Seven of the top hits (*BCKDHA*, *EFNA1*, *NEK6*, *PKLR*, *RNASET2*, *SLC1A1*, or *TLR3*) were validated in a secondary screen.

### Immunoblotting

Cells were washed ice-cold 1X PBS and then lysed on ice in lysis buffer [50 mM Tris.Cl (pH 7.5), 400 mM NaCl, 1% Nonidet P-40, 1 mM EDTA, and 10% glycerol] freshly supplemented with a PIC tablet for 30 mins at 4°C. Extracted proteins were quantified using the Bradford Assay (Biorad 5000006) and ∼30-50 μg protein was analyzed by SDS-PAGE. The following primary antibodies were used for immunoblot analysis: ARNT (Cell Signaling Technology, #5537; RRID:AB_10694232), GLUT1 (Cell Signaling Technology, #12939, RRID:AB_2687899), HIF1α (Cell Signaling Technology, #14179, RRID:AB_2622225), HIF2α (Cell Signaling Technology, #71565), NDRG1 (Cell Signaling Technology, #5196, RRID:AB_10626626), SLC1A1/EAAT3 (Cell Signaling Technology, #14501, RRID:AB_2798499), TBP (Cell Signaling Technology, #44059, RRID:AB_2799258), Tubulin (Cell Signaling Technology, #2128, RRID:AB_823664), Vinculin (Cell Signaling Technology, #13901, RRID:AB_2728768), V5 Tag (Cell Signaling Technology, #80076, RRID:AB_2920661). All antibodies were used at 1:1000 dilutions. Primary antibodies were detected with HRP-conjugated secondary antibodies (Pierce, 1:5000) and chemiluminescent HRP substrates (Supersignal West Pico PLUS, Thermo Fisher Scientific; Pierce ECL Plus, Thermo Fisher Scientific; or Immobilon ECL Ultra, Millipore) (Thermo Fisher Scientific, RRID:SCR_008452).

### Protein Modeling

Glutamate (green) was docked into the co-crystal structure of SLC1A1/EAAT3 + aspartate (PDB: 6X2Z; grey) and subsequently energy minimized and refined using MOE. 2D ligand interaction diagrams were generated in MOE, while the 3D renderings were generated in ChimeraX (h-bonds from aspartate in pink). Frequently mutated Arg445, which does not directly interact with the amino acid substrates, is highlighted in cyan.

### Metabolomics (LC-MS/MS)

Targeted Mass Spectrometry. Samples were re-suspended using 20 μl HPLC grade water for mass spectrometry. 5-7 μl were injected and analyzed using a hybrid 6500 QTRAP triple quadrupole mass spectrometer (AB/SCIEX) coupled to a Prominence UFLC HPLC system (Shimadzu) via selected reaction monitoring (SRM) of a total of 298 endogenous water-soluble metabolites for steady-state analyses of samples. Some metabolites were targeted in both positive and negative ion mode for a total of 309 SRM transitions using positive/negative ion polarity switching. ESI voltage was +4950V in positive ion mode and –4500V in negative ion mode. The dwell time was 3 ms per SRM transition and the total cycle time was 1.55 seconds. Approximately 9-12 data points were acquired per detected metabolite. Samples were delivered to the mass spectrometer via hydrophilic interaction chromatography (HILIC) using a 4.6 mm i.d x 10 cm Amide XBridge column (Waters) at 400 μL/min. Gradients were run starting from 85% buffer B (HPLC grade acetonitrile) to 42% B from 0-5 minutes; 42% B to 0% B from 5-16 minutes; 0% B was held from 16-24 minutes; 0% B to 85% B from 24-25 minutes; 85% B was held for 7 minutes to re-equilibrate the column. Buffer A was comprised of 20 mM ammonium hydroxide/20 mM ammonium acetate (pH=9.0) in 95:5 water:acetonitrile. Peak areas from the total ion current for each metabolite SRM transition were integrated using MultiQuant v3.0 software (AB/SCIEX).

### TCGA Analysis

HTSeq counts files for clear cell (TCGA-KIRC), papillary (TCGA-KIRP), and chromophobe (TCGA-KICH), renal cell carcinomas were downloaded from the Genomic Data Commons Data Portal. Counts were filtered and normalized using EdgeR. Principal component analysis was performed on RCC tumors using all genes and clustering by K-means was based on RCC subtype. ccRCC samples were stratified using either *CA9* or *SLC1A1* expression into the top 75%tile (representing the high group) and bottom 25%tile (representing the low group). The high and low groups were compared using Gene Set Enrichment Analysis to identify functional enrichment and EdgeR to determine differentially expressed genes.

### Functional enrichment analysis

GSEA software was downloaded from the Gene Set Enrichment Analysis website [http://www.broad.mit.edu/gsea/downloads.jsp]. GSEA was performed on the EdgeR normalized counts per million using the ‘Hallmark’ gene sets for identification of enriched/depleted signatures. Gene Sets with an FDR<0.10 and a nominal *p*-value of <0.05 were considered significant. Differentially expressed between SLC1A1-high versus low patients were rank ordered by fold change in expression and the top 100 upregulated genes were used as input for STRING-DB.

### γH2AX Staining and Foci Counts

UMRC-6 cells expressing SLC1A1 (WT) or SLC1A1 (RW) were seeded onto coverslips in a 6-well plate at 200,000 cells per well, attached overnight, and then treated with a 0-500 nM dose range of CB-839 (Selleck, S7655) for 48 hours. Cells were fixed using 4% formaldehyde for 10 minutes, permeabilized using 0.4% Triton-X, blocked in 5% normal goat serum (NGS) in 1X PBS, and incubated overnight at 4°C in anti-phospho-histone H2A.X (Ser139) (Millipore, 05-636-I, RRID:AB_2755003, 1:400). The following day, cover slips was washed thrice using 1% NGS in 1X PBS and then incubated with Alexa Fluor 488 conjugated secondary antibody for 45 minutes at room temperature (RT). Finally, cells were mounted in Vectashield mounting media with DAPI. Images were taken using an inverted fluorescence microscope and γH2AX foci were counted using ImageJ software. DAPI signal was used to create ROIs of each nucleus (setAutoThreshold > Find Edges > Analyze Particles), which were then overlayed onto the green channel. Each overlay was individually cropped and background subtracted (Duplicate > Subtract Background, rolling=10). Finally, foci within each ROI were counted [Find Maxima, noise=10, output=(Count)].

### Human Tumor Analysis

Human tumor data was compiled from either public datasets (e.g., TCGA and NCBI SRA) or from human renal cancer biospecimens that were collected upon consent in accordance with approved institutional IRBs (CCF IRB: 4639; MSKCC IRB: 06-107 and 12-237; DFCI/BWH IRB: 01-130). In the MSKCC cohorts, RNA-seq derived gene expression data was used to sort the tumor samples into quartiles based on SLC1A1 expression. LC-MS/MS based targeted metabolomics datasets of the two cohorts (top SLC1A1 versus bottom SLC1A1 expression quartile) were compared using Wilcoxon signed-rank tests and p-values were adjusted using Benjamini-Hochberg correction. Metabolites were mapped to pathways using KEGG annotations. Transcriptomic analysis was performed using DESeq2 (PMID 25516281) to identify differentially expressed genes.

Differential abundance (DA) scores evaluated whether a metabolic pathway was differentially abundant between the two groups. DA scores were calculated by applying a Wilcoxon rank-sum test to all metabolites in a pathway, and p-values were then corrected using the Benjamini-Hochberg correction (p < 0.05). The DA score for each pathway is calculated as:

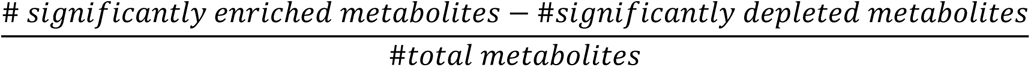

Only pathways with 3 or more significantly altered metabolites were scored.

In the DFCI/BWH cohorts, the RCC tissue microarray (TMA) analysis used two previously constructed RCC TMAs, which included tissue samples from 169 patients with localized or metastatic RCC. Each RCC case was represented by four cores taken from its tumor tissue. In addition, multiple cores representing normal kidney parenchyma and benign tumor entities were included in both TMAs.

Immunohistochemical staining of TMA sections with SLC1A1/EAAT3 antibody (Cell Signaling Technology, #14501 clone, RRID:AB_2798499, 1:100) was performed using an automated staining system (Bond III, Leica Biosystems). Semi-quantitative scoring was performed for the intensity of staining of tumor cell membranes (0 = negative; 1 = weak; 2 = moderate; 3 = strong). The H-score was then calculated as the sum of intensity scores, each multiplied by the percentage of positive tumor cells with that intensity level. The cores in the TMA were scored independently by two pathologists who subsequently reviewed the scores together to reach final consensus scores.

### Pharmacological Studies

The pharmacological properties of Cmpd <u>3e</u> were analyzed by Q2 Solutions as a fee-for-service by using turbo ion spray liquid chromatography/tandem mass spectrometry (LC/MS/MS). All samples tissue were extracted using protein precipitation in the presence of Diclofenac (100.00 ng/mL in 1:1 MeOH/ACN) used as internal standard. The parameters in the table below describe the details of the analytical pipeline:

|  |  |
| --- | --- |
| Analyte | Cmpd 3e |
| Strain / Species | CD-1 Mouse |
| Analytical Matrix /<br>Anticoagulant | Plasma: K2EDTA<br>Brain: NA<br>Kidney: NA |
| Internal Standard (IS) | Diclofenac |
| Calibration Curve Range –<br>lower limit of quantitation<br>(LLOQ) to upper limit of<br>quantitation (ULOQ) | Plasma: 2.50 ng/mL – 5,000.00 ng/mL<br>Brain: 1.00 ng/mL – 50,000.00 ng/mL<br>Kidney: 1.00 ng/mL – 50,000.00 ng/mL |
| Regression, Weighting | Quadratic, 1/ x<sup>2</sup> |
| Mass Spectrometer | AB Sciex API 4000 |
| Mobile Phase (A) | Water/1M NH<sub>4</sub>HCO<sub>3</sub> (2000:10, v/v) |
| Mobile Phase (B) | MeOH/1M NH<sub>4</sub>HCO<sub>3</sub> (2000:10, v/v) |
| Injector Wash Solvent | Acetone:Water:Formic Acid (2500:2500:50, v/v) |
| LC Pumps | Nexera LC-30AD |
| LC Controller | Shimadzu CBM-20A |
| Autosampler | PAL System |
| Column | Sprite Echelon C18 4 μm 20x2.1mm |
| Injection Volume | 10 μL |
| Transitions Monitored<br>[$\pm 0.2$ for each mass-to-charge ratio ($m/z$)] | Cmpd 3e, $m/z$ 368.2 → $m/z$ 155.9<br>Diclofenac, $m/z$ 296.0 → $m/z$ 214.0 |
| K2EDTA = dipotassium ethylenediaminetetraacetic acid<br>$m/z$ = mass-to-charge-ratio | |

## DATA AND MATERIALS AVAILABILITY

All the data necessary to evaluate the conclusions of the manuscript are provided in the paper and/or the Supplementary Materials. All ChIP-Seq and gene expression data are deposited into the Gene Expression Omnibus (Gene Expression Omnibus (GEO), RRID:SCR_005012) under accession number GSE241864. All reagents are available commercially and published constructs will be deposited in Addgene to ensure public distribution.

## Author contributions

***Conceptualized, Designed, and Analyzed Experiments*** - TG, JK, SM, FG, ND, CS, DAO, MGG, SRM, PK, JAC, CT, ER, JMA, SRS, SS, WGK, AAC; ***Performed experiments*** - TG, JK, SM, FG, ND, CS, DAO, MGG, SRM, PK, JAC, CT, AAC; ***Generated Patient Datasets*** - ER, RRK, AAK, NA, CW, TKC, SS; ***Wrote Manuscript*** – AAC.

---

**Supplementary Fig S1.**
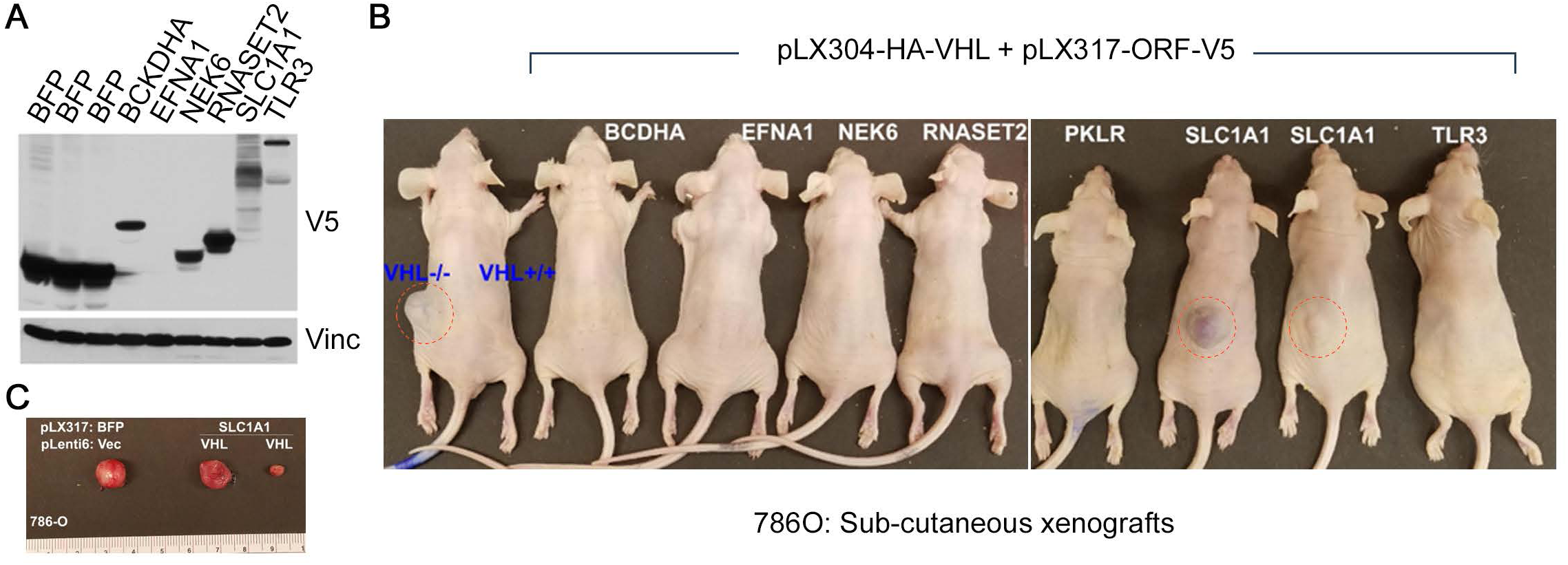
SLC1A1 Expression is Sufficient to Drive Tumor Growth. (**A** to **C**) Immunoblot analysis (**A**) and sub-cutaneous tumor growth (**B** and **C**) measured in the indicated pVHL-deficient or pVHL-proficient versions of 786-O cells lentivirally transduced to express the indicated genes. Cells were inoculated sub-cutaneously into flanks of NCR^nu/nu^ mice and tumors harvested after 12 weeks of growth.

**Supplementary Fig S2.**
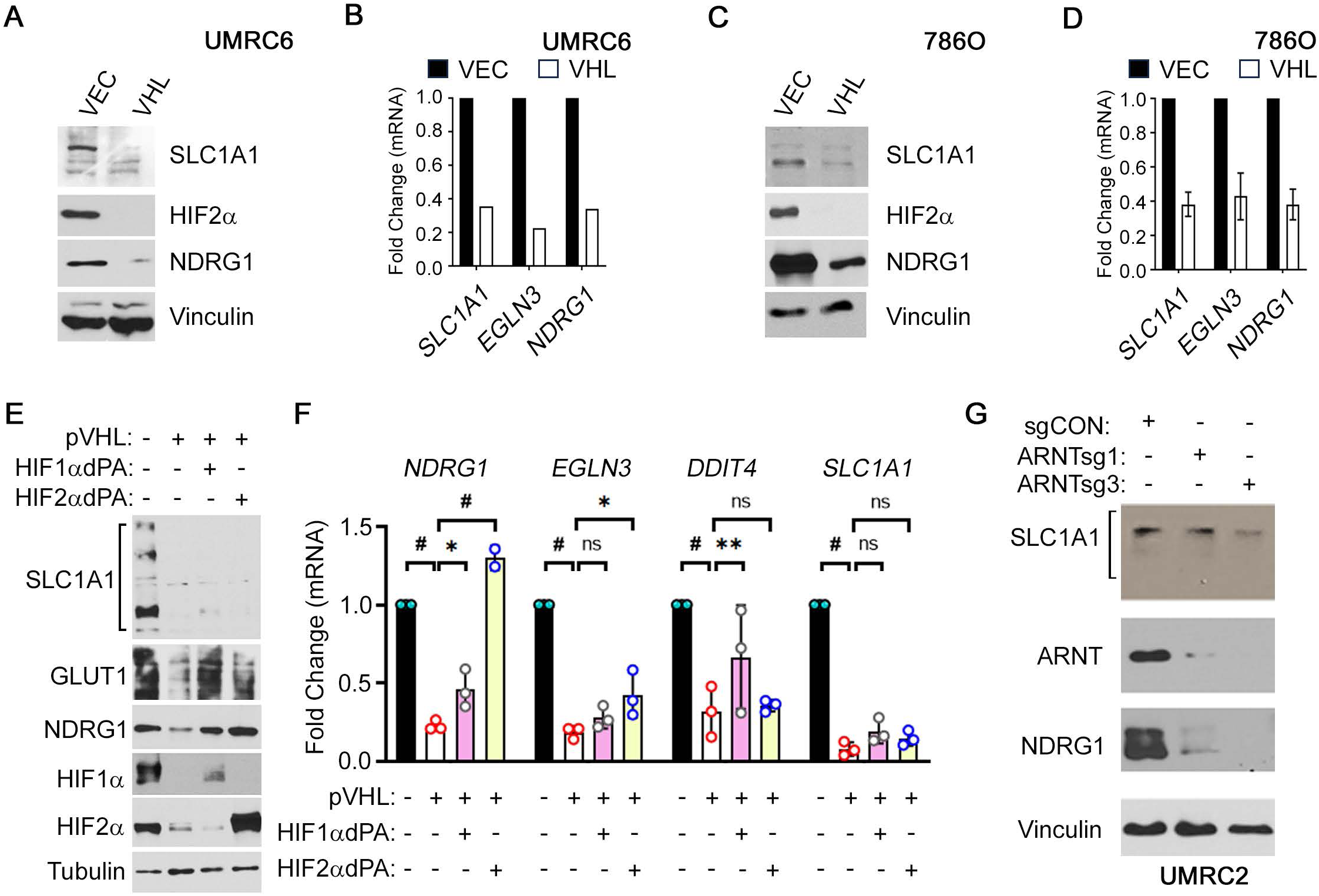
SLC1A1 Expression is pVHL Dependent. (**A** to **D**) Immunoblotting and quantitative PCR of the indicated genes in pVHL-proficient (VHL) and pVHL-deficient (VEC) versions of UMRC-6 (**A** and **B**) and 786-O (**C** and **D**) ccRCC cell lines. (**E** and **F**) Immunoblot analysis of the indicated proteins (**E**) and real-time qPCR of the indicated genes (**F**) in pVHL-proficient UMRC-2 cells lentivirally transduced to express non-degradable versions of HIF (HIF1αdPA and HIF2αdPA), or pVHL-deficient UMRC-2 cells. (**G**) Immunoblot analysis of the indicated proteins in pVHL-deficient UMRC-2 cells lentivirally transduced with CRISPR/Cas9 sgRNAs targeting *ARNT* (ARNTsg1 and ARNTsg3) or non-targeting controls (sgCON), as indicated. Expression analysis was done post selection using Puro (2 µg/ml) for 3-5 days.

**Supplementary Fig S3.**
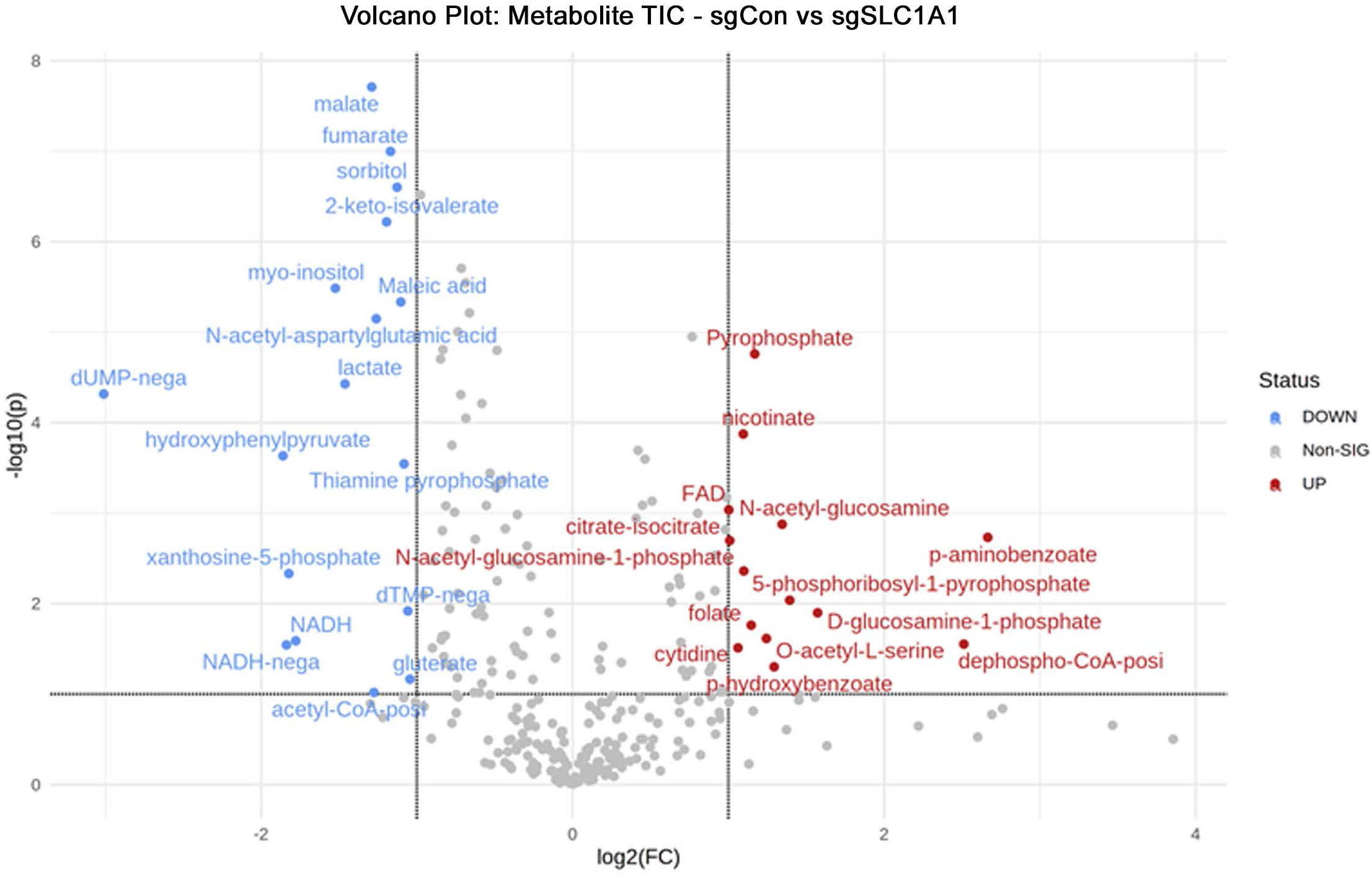
SLC1A1 Inactivation Metabolically Reprograms ccRCCs. Volcano plot comparing changes in metabolite total ion counts (TIC), as measured using LC-MS/MS analysis and analyzed using MetaboAnalyst, in pVHL-deficient UMRC-2 cells lentivirally transduced to express CRISPR/Cas9 sgRNAs that target *SLC1A1* (sgSCL1A1) or a non-targeting control (sgCon). Metabolites were harvested one day post-selection before the emergence of overt cytotoxicity. Directionality of log_2_fold-changes indicate increases (>1) or decreases (<-1) in sgSLC1A1 versus sgCon.

**Supplementary Fig S4.**
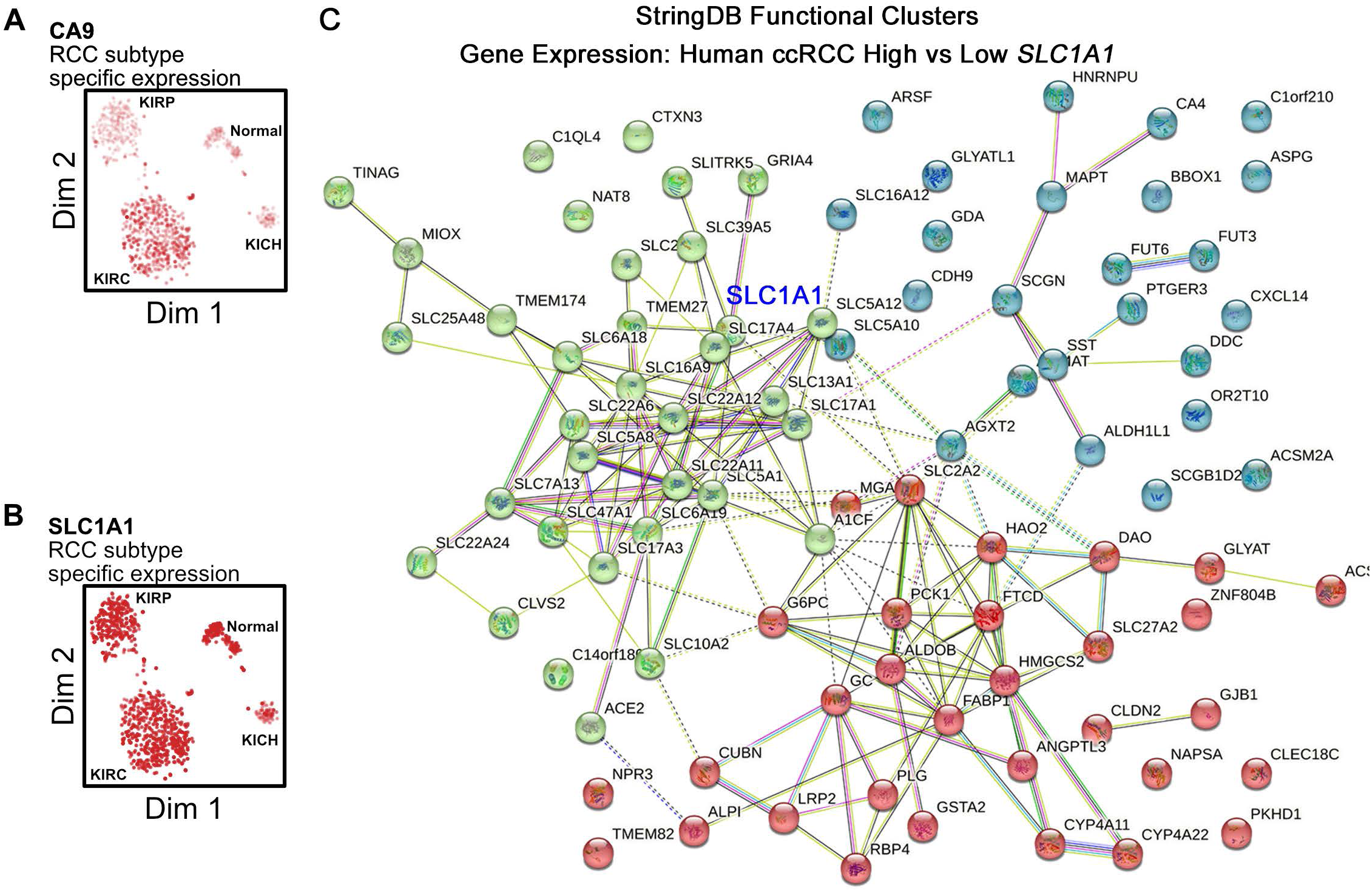
SLC1A1 Expression is Associated with Dysregulation of Other Metabolite Carriers in ccRCC. (**A** and **B**) Principal components analysis and T-Stochastic Neighbor Embedding using TCGA human renal cancer data (n=135) showing clustering of ccRCC patient samples compared to other RCC subtypes. Color overlay indicates level of expression of *CA9* (**A**) or *SLC1A1* (**B**). (**C**) StringDB analysis on the top 100 upregulated genes associated with high *SLC1A1* expression in the MSKCC cohort of ccRCC patient samples representing a clustered metabolic hub of multiple dysregulated SLCs.

**Supplementary Fig S5.**
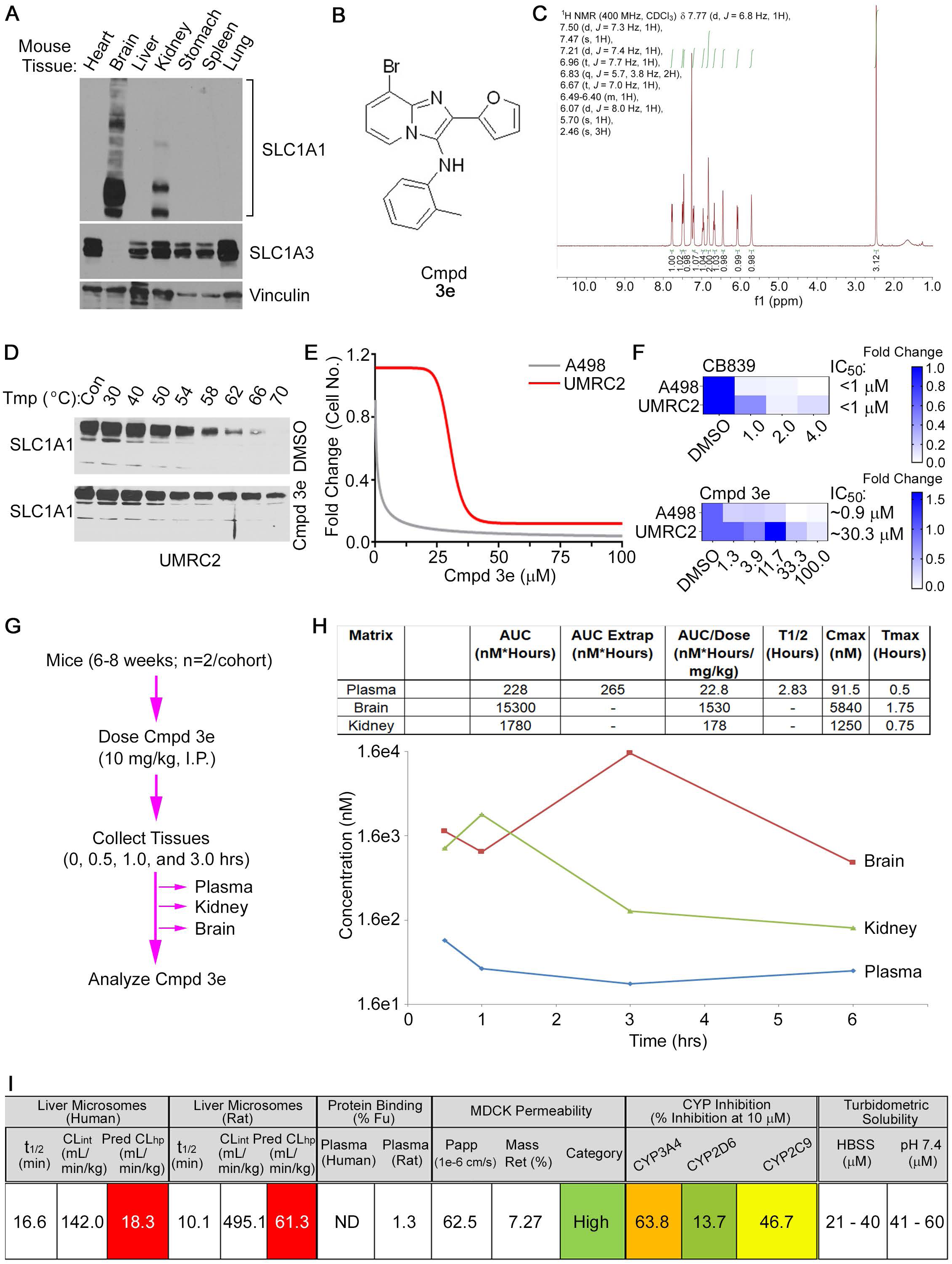
Pharmacological Blockade of SLC1A1 is Feasible In Vitro but not In Vivo. (**A**) Immunoblot analysis of the indicated proteins in the indicated normal mouse tissues. (**B** and **C**) Structure (**B**) and NMR analysis (**C**) of Cmpd <u>3e</u>. (**D**) Cellular Thermal Shift Assay (CETSA) comparing stability of SLC1A1 in whole cell extracts treated ex vivo with 100 µM Cmpd <u>3e</u> or vehicle (DMSO) and then heated to the indicated temperature for 5 mins. (**E**) Cell viability curves, plotted using Graphpad, using cell viability measured using Cell-titer Glo in two ccRCC cell lines (A-498 and UMRC-2) seeded at low density and then treated with the indicated doses of Cmpd <u>3e</u> for 7 days. (**F**) Heatmaps representing fold change in cell viability, as measured using Cell-titer Glo, in two ccRCC cells lines (A-498 and UMRC-2) that were treated with Cmpd <u>3e</u> for 7 days or CB-839 for 3 days. (**G** to **I**) Schema for pharmacological studies (**G**), concentrations of Cmpd <u>3e</u> in the indicated tissues at the indicated times post injection (**H**), and pharmacological properties of Cmpd <u>3e</u> (**I**), using tissues from mice that were dosed intraperitonially with 10 mg/kg of Cmpd <u>3e</u> or vehicle control.

**Supplementary Fig. S6.**
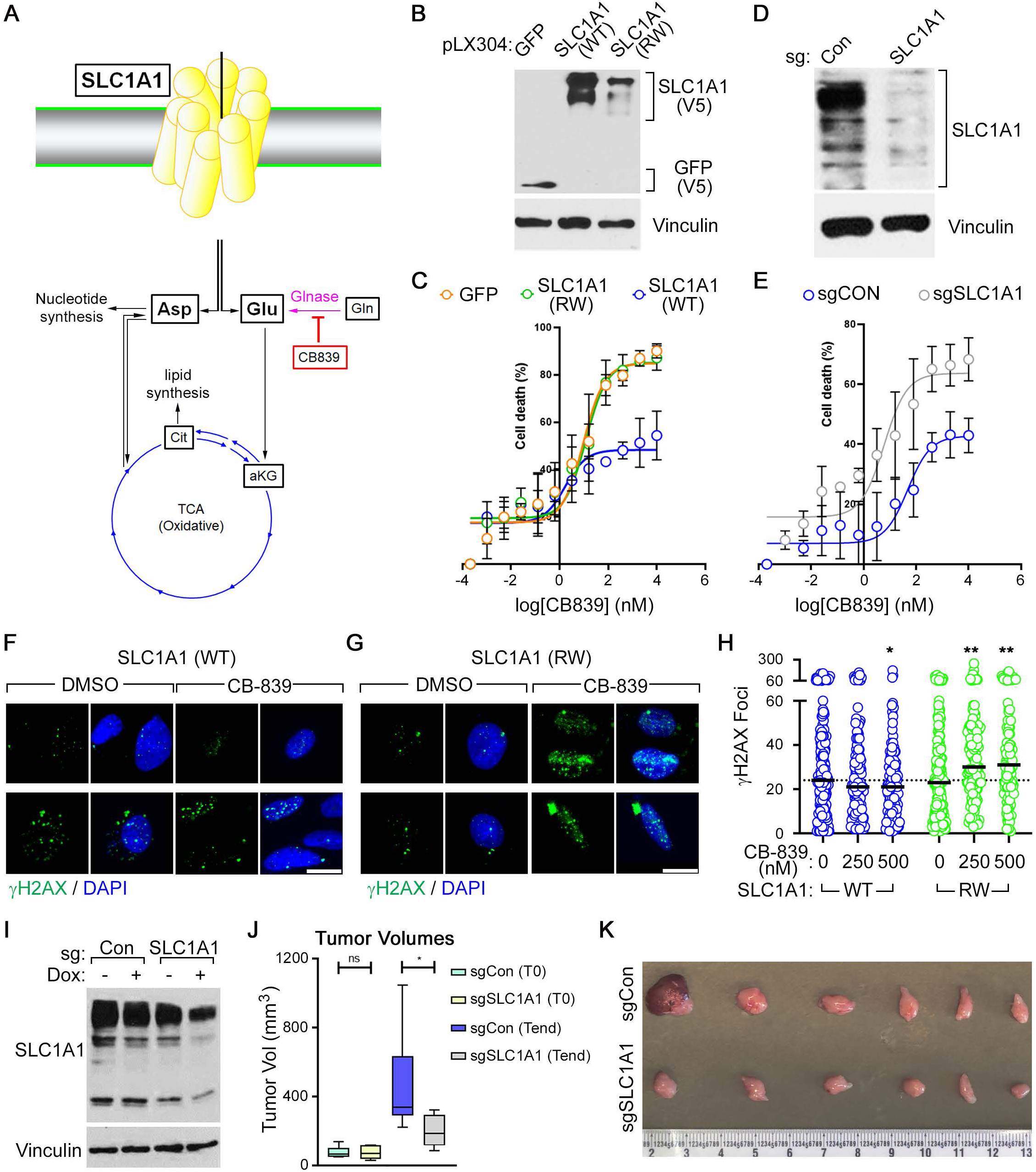
SLC1A1 Inactivation is Clinically Relevant. (**A**) Schema for the functional interaction between SLC1A1 and Glutaminase. (**B** and **C**) Immunoblot analysis (**B**) and CB-839 cytotoxicity following 3 days of treatment at the indicated concentrations, as measured using CellTiter-Glow (**C**), in UMRC-6 cells that were lentivirally transduced to express either wild-type SLC1A1 (WT), R445 mutant SLC1A1 (RW), or GFP (control). (**D** and **E**) Immunoblot analysis (**D**) and CB-839 cytotoxicity following 3 days of treatment at the indicated concentrations, as measured using CellTiter-Glow (**C**), in UMRC-2 cells that were lentivirally transduced to express sgRNAs targeting SLC1A1 or non-targeting control (sgCON). In (**C**) and (**E**), data is represented as mean±S.D. (n=3) % cell death, relative to DMSO control. (**F** to **H**) Representative immunofluorescence images of γH2AX staining in UMRC-6 cells lentivirally transduced to express either wild-type SLC1A1 (WT) (**F**) or R445 mutant SLC1A1 (RW) (**G**), treated with either 500 nM CB-839 or vehicle (DMSO) for 48 hours. (**H**) Quantification of γH2AX staining using ImageJ (n>100 per condition) in cells described in (**F**) and (**G**) treated with the indicated doses of CB-839 for 48 hours. Significance was determined using One-way Anova, *p<0.05, **p<0.001. (**I**) Immunoblot analysis of UMRC-2 cells that were lentivirally transduced to express doxycycline (Dox)-inducible versions of sgRNAs targeting SLC1A1 or non-targeting control (Con). (**J** and **K**) Tumor volumes, using cells described in (**I**), measured at the start of dox treatment (T0) versus the assay endpoint at two weeks of Dox treatment (Tend) (**J**) and photographs (**K**) of tumors that were harvested at the assay endpoint. In (**J**), tumor volumes in the sgSLC1A1 arm were compared to sgCon at the corresponding time-points using the Mann-Whitney test, n=6, *p<0.05; ns=non-significant.

**Supplementary Fig S7.**
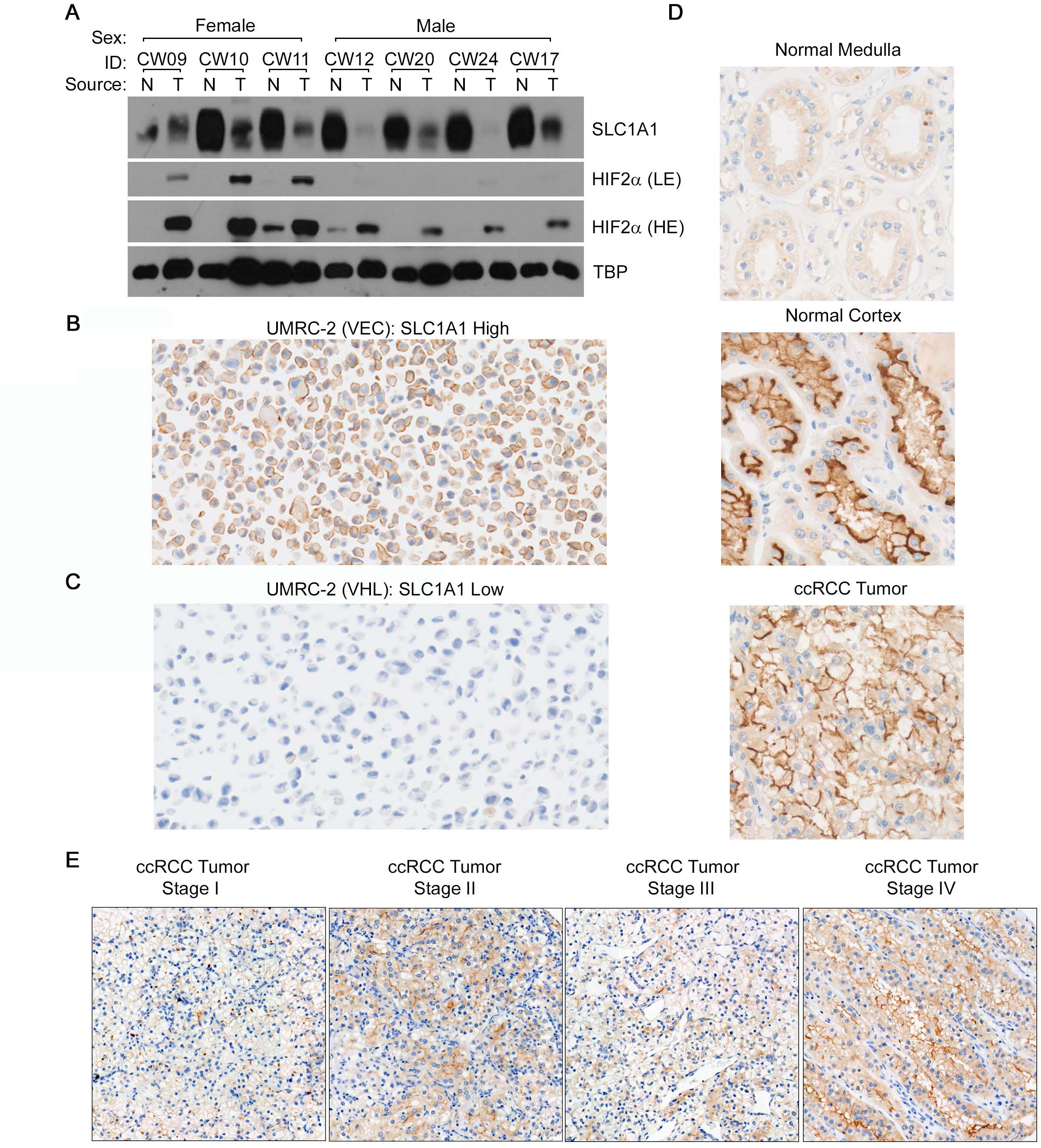
SLC1A1 Shows Dysregulated Localization in ccRCC and Correlates with Tumor Staging. (**A**) Immunoblot analysis of the indicated proteins using bulk protein lysates of ccRCC tumors (T) or paired normal adjacent tissue (N) from kidney cancer patients undergoing nephrectomies at the Cleveland Clinic. (**B** and **C**) Photomicrographs of SLC1A1 expression performed using FFPE sections of either pVHL-deficient (SLC1A1-high) (**B**) or pVHL-proficient (SLC1A1-low) (**C**) UMRC-2 cells. (**D**) Representative photomicrographs showing dysregulated sub-cellular localization of SLC1A1 in tumor versus normal kidney. Note the lack of discernible expression in the normal medulla, apical expression in the normal cortex, and expression on the entire cell surface in ccRCC tumors. (**E**) Representative photomicrographs showing SLC1A1 levels in tumors representing the indicated ccRCC stages.

## REFERENCES

1. Siegel, R.L., et al., Cancer statistics, 2022. CA Cancer J Clin, 2022. 72(1): p. 7–33.

2. Ricketts, C.J., et al., The Cancer Genome Atlas Comprehensive Molecular Characterization of Renal Cell Carcinoma. Cell Rep, 2018. 23(1): p. 313–326 e5.

3. Kaelin, W.G., Jr., The von Hippel-Lindau tumour suppressor protein: O2 sensing and cancer. Nat Rev Cancer, 2008. 8(11): p. 865–73.

4. Ivan, M., et al., HIFalpha targeted for VHL-mediated destruction by proline hydroxylation: implications for O2 sensing. Science, 2001. 292(5516): p. 464–8.

5. Jaakkola, P., et al., Targeting of HIF-alpha to the von Hippel-Lindau ubiquitylation complex by O2-regulated prolyl hydroxylation. Science, 2001. 292(5516): p. 468–72.

6. Kondo, K., et al., Inhibition of HIF2alpha is sufficient to suppress pVHL-defective tumor growth. PLoS Biol, 2003. 1(3): p. E83.

7. Gordan, J.D., et al., HIF-2alpha promotes hypoxic cell proliferation by enhancing c-myc transcriptional activity. Cancer Cell, 2007. 11(4): p. 335–47.

8. Cho, H., et al., On-target efficacy of a HIF-2alpha antagonist in preclinical kidney cancer models. Nature, 2016. 539(7627): p. 107–111.

9. Chen, W., et al., Targeting renal cell carcinoma with a HIF-2 antagonist. Nature, 2016. 539(7627): p. 112–117.

10. Jonasch, E., et al., Belzutifan for Renal Cell Carcinoma in von Hippel-Lindau Disease. N Engl J Med, 2021. 385(22): p. 2036–2046.

11. Turajlic, S., et al., Deterministic Evolutionary Trajectories Influence Primary Tumor Growth: TRACERx Renal. Cell, 2018. 173(3): p. 595–610 e11.

12. Turajlic, S., et al., Tracking Cancer Evolution Reveals Constrained Routes to Metastases: TRACERx Renal. Cell, 2018. 173(3): p. 581–594 e12.

13. Liao, L., J.R. Testa, and H. Yang, The roles of chromatin-remodelers and epigenetic modifiers in kidney cancer. Cancer Genet, 2015. 208(5): p. 206–14.

14. Xia, X., et al., Integrative analysis of HIF binding and transactivation reveals its role in maintaining histone methylation homeostasis. Proc Natl Acad Sci U S A, 2009. 106(11): p. 4260–5.

15. Varela, I., et al., Exome sequencing identifies frequent mutation of the SWI/SNF complex gene PBRM1 in renal carcinoma. Nature, 2011. 469(7331): p. 539–42.

16. Dalgliesh, G.L., et al., Systematic sequencing of renal carcinoma reveals inactivation of histone modifying genes. Nature, 2010. 463(7279): p. 360–3.

17. Chakraborty, A.A., et al., HIF activation causes synthetic lethality between the VHL tumor suppressor and the EZH1 histone methyltransferase. Sci Transl Med, 2017. 9(398).

18. Loven, J., et al., Selective inhibition of tumor oncogenes by disruption of super-enhancers. Cell, 2013. 153(2): p. 320–34.

19. Whyte, W.A., et al., Master transcription factors and mediator establish super-enhancers at key cell identity genes. Cell, 2013. 153(2): p. 307–19.

20. Huo, F.C., et al., SHMT2 promotes the tumorigenesis of renal cell carcinoma by regulating the m6A modification of PPAT. Genomics, 2022. 114(4): p. 110424.

21. Liu, Z., et al., Serine hydroxymethyltransferase 2 knockdown induces apoptosis in ccRCC by causing lysosomal membrane permeabilization via metabolic reprogramming. Cell Death Dis, 2023. 14(2): p. 144.

22. Zhang, J., et al., VHL substrate transcription factor ZHX2 as an oncogenic driver in clear cell renal cell carcinoma. Science, 2018. 361(6399): p. 290–295.

23. Yao, X., et al., VHL Deficiency Drives Enhancer Activation of Oncogenes in Clear Cell Renal Cell Carcinoma. Cancer Discov, 2017. 7(11): p. 1284–1305.

24. Chen, Y. and R.A. Swanson, The glutamate transporters EAAT2 and EAAT3 mediate cysteine uptake in cortical neuron cultures. J Neurochem, 2003. 84(6): p. 1332–9.

25. O’Kane, R.L., et al., Na(+)-dependent glutamate transporters (EAAT1, EAAT2, and EAAT3) of the blood-brain barrier. A mechanism for glutamate removal. J Biol Chem, 1999. 274(45): p. 31891–5.

26. Qiu, B. and O. Boudker, Symport and antiport mechanisms of human glutamate transporters. Nat Commun, 2023. 14(1): p. 2579.

27. Bailey, C.G., et al., Loss-of-function mutations in the glutamate transporter SLC1A1 cause human dicarboxylic aminoaciduria. J Clin Invest, 2011. 121(1): p. 446–53.

28. Kaelin, W.G., Jr. and P.J. Ratcliffe, Oxygen sensing by metazoans: the central role of the HIF hydroxylase pathway. Mol Cell, 2008. 30(4): p. 393–402.

29. Yuan, M., et al., Ex vivo and in vivo stable isotope labelling of central carbon metabolism and related pathways with analysis by LC-MS/MS. Nat Protoc, 2019. 14(2): p. 313–330.

30. Xia, J., et al., MetaboAnalyst 2.0--a comprehensive server for metabolomic data analysis. Nucleic Acids Res, 2012. 40(Web Server issue): p. W127–33.

31. Subramanian, A., et al., Gene set enrichment analysis: a knowledge-based approach for interpreting genome-wide expression profiles. Proc Natl Acad Sci U S A, 2005. 102(43): p. 15545–50.

32. Peghini, P., J. Janzen, and W. Stoffel, Glutamate transporter EAAC-1-deficient mice develop dicarboxylic aminoaciduria and behavioral abnormalities but no neurodegeneration. EMBO J, 1997. 16(13): p. 3822–32.

33. Bunch, L., M.N. Erichsen, and A.A. Jensen, Excitatory amino acid transporters as potential drug targets. Expert Opin Ther Targets, 2009. 13(6): p. 719–31.

34. Nieoullon, A., et al., The neuronal excitatory amino acid transporter EAAC1/EAAT3: does it represent a major actor at the brain excitatory synapse? J Neurochem, 2006. 98(4): p. 1007–18.

35. Bridges, R.J. and C.S. Esslinger, The excitatory amino acid transporters: pharmacological insights on substrate and inhibitor specificity of the EAAT subtypes. Pharmacol Ther, 2005. 107(3): p. 271–85.

36. Esslinger, C.S., et al., The substituted aspartate analogue L-beta-threo-benzyl-aspartate preferentially inhibits the neuronal excitatory amino acid transporter EAAT3. Neuropharmacology, 2005. 49(6): p. 850–61.

37. Liu, N., A.A. Jensen, and L. Bunch, *beta-Indolyloxy Functionalized Aspartate Analogs as Inhibitors of the Excitatory Amino Acid Transporters (EAATs)*. ACS Med Chem Lett, 2020. 11(11): p. 2212–2220.

38. Denton, T., et al., Synthesis and preliminary evaluation of trans-3,4-conformationally-restricted glutamate and pyroglutamate analogues as novel EAAT2 inhibitors. Bioorg Med Chem Lett, 2002. 12(21): p. 3209–13.

39. Shimamoto, K., et al., Syntheses of optically pure beta-hydroxyaspartate derivatives as glutamate transporter blockers. Bioorg Med Chem Lett, 2000. 10(21): p. 2407–10.

40. Wu, P., et al., Identification and Structure-Activity Relationship Study of Imidazo[1,2-a]pyridine-3-amines as First Selective Inhibitors of Excitatory Amino Acid Transporter Subtype 3 (EAAT3). ACS Chem Neurosci, 2019. 10(10): p. 4414–4429.

41. Gameiro, P.A., et al., In vivo HIF-mediated reductive carboxylation is regulated by citrate levels and sensitizes VHL-deficient cells to glutamine deprivation. Cell Metab, 2013. 17(3): p. 372–85.

42. Jiang, L., et al., Reductive carboxylation supports redox homeostasis during anchorage-independent growth. Nature, 2016. 532(7598): p. 255–8.

43. Metallo, C.M., et al., Reductive glutamine metabolism by IDH1 mediates lipogenesis under hypoxia. Nature, 2011. 481(7381): p. 380–4.

44. Okazaki, A., et al., Glutaminase and poly(ADP-ribose) polymerase inhibitors suppress pyrimidine synthesis and VHL-deficient renal cancers. J Clin Invest, 2017. 127(5): p. 1631–1645.

45. Guo, W., et al., Dysregulated Glutamate Transporter SLC1A1 Propels Cystine Uptake via Xc(-) for Glutathione Synthesis in Lung Cancer. Cancer Res, 2021. 81(3): p. 552–566.

46. Xiong, J., et al., SLC1A1 mediated glutamine addiction and contributed to natural killer T-cell lymphoma progression with immunotherapeutic potential. EBioMedicine, 2021. 72: p. 103614.

47. Wang, X., et al., SLC1A1-mediated cellular and mitochondrial influx of R-2-hydroxyglutarate in vascular endothelial cells promotes tumor angiogenesis in IDH1-mutant solid tumors. Cell Res, 2022. 32(7): p. 638–658.

48. Garcia-Bermudez, J., et al., Aspartate is a limiting metabolite for cancer cell proliferation under hypoxia and in tumours. Nat Cell Biol, 2018. 20(7): p. 775–781.

49. Hakimi, A.A., et al., An Integrated Metabolic Atlas of Clear Cell Renal Cell Carcinoma. Cancer Cell, 2016. 29(1): p. 104–116.

50. Orlando, D.A., et al., Quantitative ChIP-Seq normalization reveals global modulation of the epigenome. Cell Rep, 2014. 9(3): p. 1163–70.

